# Delta opioid receptors engage multiple signaling cascades to differentially modulate prefrontal GABA release with input and target specificity

**DOI:** 10.1101/2024.08.08.607246

**Authors:** Ryan P. D. Alexander, Kevin J. Bender

**Author notes:** Correspondence (R.P.D.A.), (K.J.B.).

## Abstract

Opioids regulate circuits associated with motivation and reward across the brain. Of the opioid receptor types, delta opioid receptors (DORs) appear to have a unique role in regulating the activity of circuits related to reward without a liability for abuse. In neocortex, DORs are expressed primarily in interneurons, including parvalbumin- and somatostatin-expressing interneurons that inhibit somatic and dendritic compartments of excitatory pyramidal cells, respectively. But how DORs regulate transmission from these key interneuron classes is unclear. We found that DORs regulate inhibition from these interneuron classes using different G-protein signaling pathways that both converge on presynaptic calcium channels, but regulate distinct aspects of calcium channel function. This imposes different temporal filtering effects, via short-term plasticity, that depend on how calcium channels are regulated. Thus, DORs engage differential signaling cascades to regulate inhibition depending on the postsynaptic target compartment, with different effects on synaptic information transfer in somatic and dendritic domains.

## INTRODUCTION

The opioid receptor family is comprised of three isoforms—mu, delta, and kappa—that are expressed throughout cortical and subcortical brain regions^1-3^. Although mu receptors are responsible for the main analgesic and addictive effects of opioid painkillers and narcotics^4,5^, delta opioid receptors (DORs) play a critical modulatory role in pain and reward circuitry^6-8^. While not habit-forming on their own^7^, DORs contribute strongly to the development of reward associations. Indeed, DOR antagonists reduce conditioned place preference (CPP) for addictive drugs including morphine^9-11^, even though DORs cannot bind morphine themselves^12^. DORs are enriched in anterior cortical areas like the medial prefrontal cortex (mPFC)^13-15^, where dysregulated opioidergic signaling is associated with impulsivity and drug-seeking behavior^16,17^. Remarkably, selective DOR knockdown in specific interneuron subpopulations in mPFC can prevent morphine-induced CPP^18^. Though DORs in mPFC, particularly in GABAergic interneurons, appear to have central importance in reward processing, how DORs regulate interneuron function remains unclear.

DORs primarily influence neuronal activity by modulating transmitter release from presynaptic terminals^19,20^. Canonically, DORs couple to G_i/o_ signaling cascades^21^. Following receptor activation, Gβγ subunits dissociate from DORs and inhibit presynaptic voltage-gated calcium channels (Ca_V_)^22^ by depolarizing the voltage-dependence of channel activation^23-25^. This leads to a net reduction in intracellular Ca required for transmitter release, and thus a reduction in vesicle release probability (*P*_*R*_)^26,27^. This style of neuromodulation is common across multiple G_i/o_-coupled receptors, including presynaptic metabotropic glutamate, GABA_B_, endocannabinoid, and catecholamine receptors^28,29^. Reductions in *P R* typically increase the relative amplitude of subsequent events, a process termed short-term synaptic facilitation. This form of short-term plasticity (STP) is due to complex temporal dynamics of Ca in presynaptic boutons, Ca buffering mechanisms, and vesicle release proteins^30^. Presynaptic inhibition by G-protein signaling is ubiquitous throughout the brain and mostly results in increased STP^28,29^. However, at a variety of synapses, DORs and other opioid receptors appear to break this rule, inhibiting release with little to no increase in STP^31-34^. Why this occurs has remained largely unexplained.

Here, we studied DOR modulation in two subtypes of GABAergic interneurons that inhibit layer 5 pyramidal cells: parvalbumin-expressing (PV+) interneurons, which target the perisomatic regions, and somatostatin-expressing (SOM+) interneurons, which target dendritic regions, including apical dendrites that span the upper layers of cortex. We found that DORs suppressed GABA release in PV+ cells via well-studied, canonical signaling pathways where Gβγ subunits alter the voltage-dependence of activation of presynaptic Ca_V_s, thereby reducing *P*_*R*_ and increasing STP. By contrast, SOM+ cell transmission was regulated via multiple DOR-dependent signaling cascades, engaged in parallel in the same bouton. DORs engaged both canonical Gβγ-dependent modulation of Ca_V_s and a second, non-canonical pathway that was completely independent of G_i/o_-based signaling. This second pathway also regulated presynaptic Ca_V_s, but through a reduction in *P*_*R*_ without increasing STP—a mechanism recently described for dopaminergic regulation of glutamatergic transmission in mPFC^35^. These different forms of regulation at PV+ and SOM+ terminals thus produced differential temporal filtering of inhibition that varied depending on inhibitory cell target. Taken together, these results show that DORs regulate inhibitory transmission in mPFC through multiple presynaptic mechanisms, even within a single axonal bouton of SOM+ interneurons. Further, this demonstrates that neuromodulators can engage multiple signaling cascades simultaneously to impose different temporal filtering rules depending on target structure.

## RESULTS

### Unconventional regulation of prefrontal GABA release by delta opioid receptors

DORs are highly expressed by GABAergic interneurons in mPFC but how they regulate inhibitory transmission is not known. To test this, we made whole-cell recordings from layer 5 (L5) pyramidal cells in slices containing mPFC and evoked inhibitory postsynaptic currents (eIPSCs) with a bipolar stimulating electrode (20 Hz, 15 s interval) (**Figure 1A**). Application of the selective DOR agonist DPDPE (1 μM) reduced eIPSC amplitude (Normalized Amplitude = 0.47 ± 0.05, n/N = 13/6; p < 0.0001, paired t-test; **Figure 1B-D**). These effects were blocked by pre-treating slices with the DOR antagonist naltrindole (2 μM; Norm Amp: 0.99 ± 0.02, n/N = 6/2; p = 0.55, paired t-test; **Figure 1C**). Deltorphin-II (1 μM), another agonist that displays preferential activation of type 2 DORs^36^, had comparable effects (Norm Amp = 0.41 ± 0.06, n/N = 9/3; p < 0.0001, paired t-test), confirming that eIPSC reduction was mediated through DORs.

**Figure 1:**
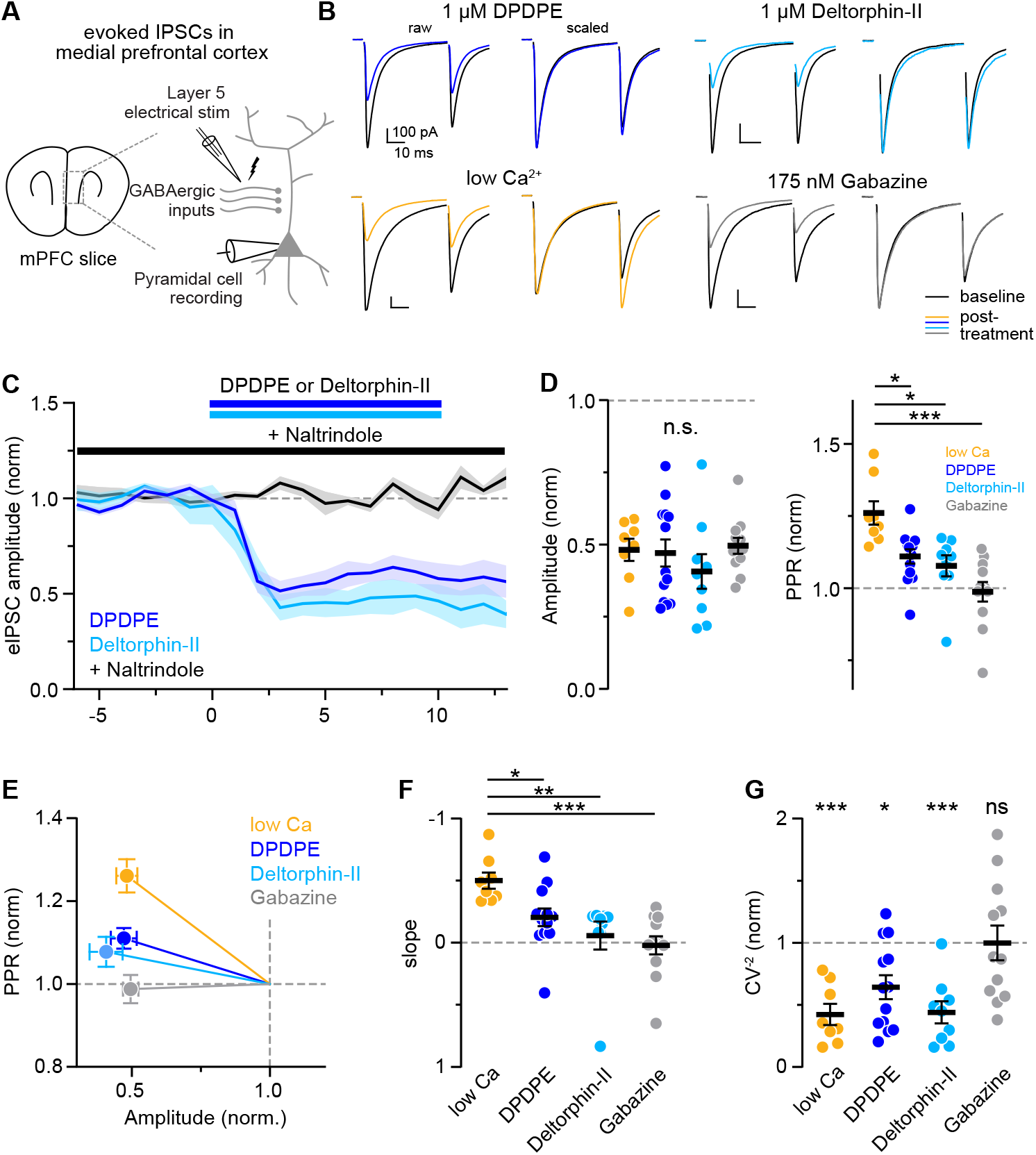
DOR signaling suppresses inhibitory transmission onto L5 pyramidal neurons in mPFC. **A:** Schematic of experimental paradigm. **B:** Example current traces evoked by paired-pulse synaptic stimulation before and after application of DPDPE (1 μM), deltorphin-II (1 μM), gabazine (175 nM), or low external Ca (0.65 mM). Synaptic currents are overlaid as either raw values (left) or normalized to the first response (right) to visualize PPR differences. Scale bars depict 100 pA and 10 ms in all conditions. **C:** Summary plot of normalized eIPSC amplitude over time during DOR agonist application. Bars depict when drug was applied. **D:** Summary plots of normalized eIPSC amplitude (left) and PPR (right) for all conditions. Colored circles represent individual cells, black bars depict mean values. **E:** Plot of normalized PPR as a function of normalized eIPSC amplitude after each treatment, with baseline at (1,1). Slope of each line represents degree of PPR change relative to degree of amplitude change. **F:** Summary plot of slope values for all conditions. **G:** Summary plot of normalized CV^-2^ for all conditions. * p<0.05, *** p<0.001.

DORs, as with other opioid receptors, are commonly located presynaptically and reduce *P*_*R*_ when activated^9,27,37-39^. Reduced *P*_*R*_ increases variability in response amplitude, as measured by coefficient of variance (CV), and typically increases short-term facilitation, as measured by paired-pulse ratio (PPR). PPR increased following DPDPE (Norm PPR = 1.11 ± 0.02; p = 0.0024, Wilcoxon signed rank test; **Figure 1D**), although the change was relatively modest (11% PPR increase vs. 53% amplitude decrease), whereas no significant change was observed with deltorphin-II (Norm PPR = 1.08 ± 0.04; p = 0.13, Wilcoxon signed rank test). Both DOR agonists reduced CV^-2^ (Norm CV^-2^; DPDPE = 0.64 ± 0.10; p = 0.0031; deltorphin-II = 0.44 ± 0.09; p = 0.0002, paired t-test; **Figure 1G**), indicating that DORs act presynaptically to decrease *P*_*R*_. In agreement with this, DPDPE had no effect on postsynaptic membrane properties in these recording conditions where potassium channels are blocked by intracellular cesium (control vs. DPDPE; ΔR_input_ = 1.4 ± 3.5 % vs. 1.3 ± 1.2 %, n/N = 5/2 vs. 29/15; p = 0.71; ΔI_hold_ = -7.7 ± 3.7 pA vs. -5.1 ± 2.2 pA; p = 0.82, Holm-Sidak post-hoc test; **Extended Data 1B-C)**. In separate experiments using a K^+^-based intracellular solution that does not block potassium channels, DPDPE modestly affected R_input_ (ΔR_input_ = -11.2 ± 2.4%, n/N = 8/3; p = 0.016; ΔI_hold_ = -13.2 ± 3.1 pA; p = 0.55, Holm-Sidak post-hoc test), suggesting that DOR may have additional postsynaptic actions under physiological conditions.

Changes in *P*_*R*_ often cause proportional reductions in response amplitude and increases in PPR^40-42^. By contrast, we observed what appeared to be relatively small increases in PPR with DOR modulation. To determine whether this reflects unique release properties of mPFC GABAergic synapses, or instead suggests that DORs modulate release without typical changes in STP, we benchmarked DOR modulation against manipulations that canonically affect *P*_*R*_ or postsynaptic components of transmission. First, external Ca concentration was reduced from 1.3 to 0.65 mM (“low Ca”), as this is known to reduce *P*_*R*_. This reduced eIPSC amplitude and CV^-2^, while causing increased PPR (Norm Amp = 0.48 ± 0.04, n/N = 8/2; p < 0.0001, paired t-test; Norm PPR = 1.26 ± 0.04; p = 0.0078, Wilcoxon signed rank test; Norm CV^-2^ = 0.42 ± 0.08; p = 0.0003, paired t-test; **Figure 1D, G**). To compare this to a purely postsynaptic form of neuromodulation, we blocked a fraction of GABA_A_ receptors with gabazine (175 nM). This depressed amplitude without altering PPR or CV^-2^ (Norm Amp = 0.50 ± 0.03, n/N = 12/3; p < 0.0001, paired t-test; Norm PPR = 0.99 ± 0.03; p = 0.91, Wilcoxon signed rank test; Norm CV^-2^ = 1.00 ± 0.14; p = 0.99, paired t-test; **Figure 1D, G**). Of note, both manipulations were tuned to produce similar reductions in IPSC amplitude, allowing for direct comparison to DOR-dependent modulation (F_3,38_ = 0.72, p = 0.55, one-way ANOVA; **Figure 1D**). When compared to these benchmarks, we found that DPDPE and deltorphin-II both resulted in less PPR change than expected based on changing external Ca concentration (DPDPE: p = 0.038; deltorphin-II: p = 0.017, Kruskal-Wallis test; **Figure 1D**), but clearly altered CV^-2^, in sharp contrast to gabazine. To better visualize these differences, normalized amplitude and PPR in each condition were plotted as X-Y coordinates with baseline values at (1.0, 1.0). This allowed us to quantify each amplitude-PPR relationship as a slope. Within this scheme, purely postsynaptic manipulations (e.g., gabazine) do not deviate from the X-axis. By contrast, canonical forms of presynaptic modulation (e.g., low Ca) are depicted with a steep inverse relationship (**Figure 1E**). Surprisingly, both DPDPE and deltorphin-II exhibited slopes that were intermediate between both of these benchmarks (e.g., low Ca vs. DPDPE: -0.50 ± 0.07 vs. -0.20 ± 0.07; p = 0.038; low Ca vs. deltorphin-II: -0.50 ± 0.07 vs. -0.06 ± 0.11; p = 0.027, Dunnett’s test; **Figure 1E-F**). These observations, coupled with clear changes in CV^-2^, suggest that DORs engage presynaptic signaling mechanisms that are difficult to explain at first sight.

### Differential DOR modulation of PV+ and SOM+ inputs

Electrical stimulation recruits GABAergic synapses independent of their source. Thus, one explanation for the mixed effects observed above is that DORs differentially filter transmission from different GABAergic inputs, and the average of this is sampled with electrical stimulation of tissue. In neocortex, DOR-encoding *Oprd1* mRNA is expressed largely in parvalbumin- (PV+) and somatostatin-expressing (SOM+) cells^43,44^ (**Figure 2A**). Therefore, we focused on release from these two cell classes. To test whether there is differential DOR regulation between PV+ and SOM+ afferents, we expressed channelrhodopsin-2 (ChR2) in either class using Cre-inducible vectors in PV- or SOM-Cre transgenic mice, respectively. Acute mPFC slices were taken 4-6 weeks post-injection and optically-evoked IPSCs (oIPSCs) were recorded in L5 pyramidal neurons (**Figure 2B**). Consistent with electrical stimulation and predictions from mRNA expression, application of DPDPE suppressed both PV- (Norm Amp = 0.33 ± 0.04, n/N = 12/5; p < 0.0001, paired t-test) and SOM-derived oIPSCs (Norm Amp = 0.43 ± 0.05, n/N = 10/4; p < 0.0001, paired t-test; **Figure 2C-D**). Although DPDPE increased PPR in both subtypes (Norm PPR; PV = 1.32 ± 0.03; p < 0.0001; SOM = 1.08 ± 0.02, p = 0.003; paired t-test), this change was far more modest for SOM-compared to PV-oIPSCs (p < 0.0001, unpaired t-test; **Figure 2D**). The amplitude-PPR slope for PV-oIPSCs was identical to that observed in low extracellular Ca (PV vs. low Ca = -0.50 ± 0.06 vs. -0.50 ± 0.07; p = 0.98, Holm-Sidak post-hoc test; **Figure 2E-F**). By contrast, the SOM-oIPSC amplitude-PPR slope was shallower (SOM vs. low Ca = -0.14 ± 0.03 vs. -0.50 ± 0.07; p = 0.0026; SOM vs. PV = -0.14 ± 0.03 vs. -0.50 ± 0.06, p = 0.0008, Holm-Sidak post-hoc test; **Figure 2F**) and mirrored that observed with electrical stimulation (SOM vs. eStim: -0.14 ± 0.03 vs. -0.20 ± 0.07, p = 0.71; Holm Sidak post-hoc test). Taken together, these data suggest that DORs engage canonical signaling mechanisms at PV+ inputs whereas effects observed at SOM+ inputs remain unexplained.

**Figure 2:**
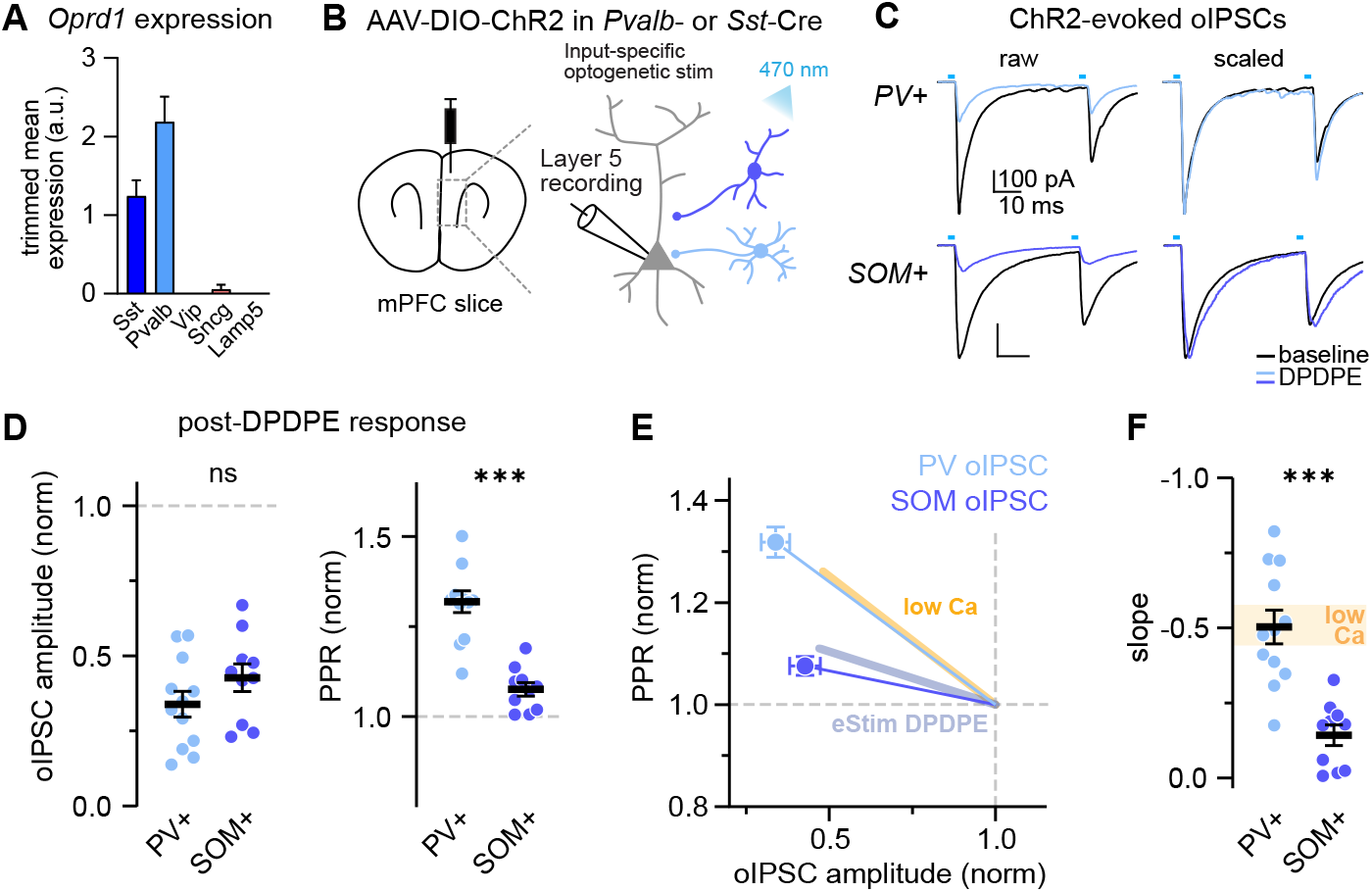
GABA release from PV+ and SOM+ interneurons is differentially regulated by DORs. **A:** Expression of *Oprd1* mRNA across GABAergic cell classes from Allen Institute mouse patch-seq database^43^. Trimmed mean expression values are averaged over subclusters of each class. **B:** Schematic depicting cell type-specific optogenetic stimulation paradigm. **C:** Example current traces of PV- (top) and SOM-derived (bottom) synaptic responses evoked by paired light pulses before and after DPDPE. **D:** Summary plots of normalized oIPSC amplitude (left) and PPR (right) between cell types. **E:** Normalized amplitude-PPR plot depicting slope relationships for PV- and SOM-oIPSCs after DPDPE, as well as for eIPSCs after DPDPE (blue line) and low Ca (yellow line) from Figure 1. **F:** Summary plot of amplitude-PPR slopes for PV- and SOM-oIPSCs after DPDPE. Yellow bar depicts low Ca eIPSC slope ± SEM for comparison. *** p<0.001.

Recently, we described a form of presynaptic neuromodulation where *P*_*R*_ is reduced without altering STP. At select excitatory inputs to mPFC pyramidal cells, activation of G_s_-coupled dopamine receptors (D1R or D5R, referred to collectively as D1R hereafter) suppressed glutamate release (i.e., lowered *P*_*R*_) without an accompanying PPR increase^35^. This was termed “gain modulation” and was mediated by a form of PKA-dependent presynaptic Ca_V_ modulation that suppressed the probability of channel opening in response to an action potential (AP). When plotted as an amplitude-PPR slope, gain modulation resembles a postsynaptic effect and lies along the X-axis. SOM+ DOR modulation is intermediate between both canonical and gain modulation forms of presynaptic regulation, suggesting that DORs do not utilize either mechanism exclusively. Therefore, we hypothesized that both processes are engaged in parallel in SOM+ terminals.

One prerequisite for this hypothesis, and gain modulation more broadly, is that Ca_V_s must be coupled to release machinery in a “nanodomain” configuration where Ca influx from an individual Ca_V_ is sufficient to trigger vesicular fusion. This configuration is common; most mature GABAergic synapses in hippocampus and cerebellum operate in a nanodomain configuration^45,46^. But whether release occurs via nanodomains in mPFC SOM+ and PV+ terminals remains unknown. This can be tested simply with divalent Ca_V_ inhibitors. Manganese (Mn) and cadmium (Cd) are both divalents that block Ca permeation but do so with different dissociation rates. Mn dissociates quickly, and mimics Gβγ-dependent canonical modulation by blocking and unblocking repeatedly during the duration of a single AP. By contrast, Cd dissociates slowly, blocking a single Ca_V_ completely for the duration of a single AP^47^. Thus, Cd will mimic gain modulation if Ca_V_ s are coupled to release machinery in a nanodomain configuration.

To test coupling configuration of mPFC interneurons, we applied external Mn or Cd and monitored L5 eIPSCs (**Extended Data 2A-D**). Both divalents supressed eIPSC amplitude and CV^-2^ dose-dependently (Norm Amp; 200 μM Mn = 0.45 ± 0.03, n/N = 11/4; p < 0.0001; 5 μM Cd = 0.56 ± 0.06, n/N = 7/3; p = 0.0004; Norm CV^-2^; Mn = 0.54 ± 0.07; p = 0.0009; Cd = 0.51 ± 0.15; p = 0.016, paired t-test), but only Mn increased PPR (Norm PPR; Mn = 1.24 ± 0.05; p = 0.001, Wilcoxon signed rank test; Cd = 0.97 ± 0.04; p = 0.52, paired t-test). Similar results were observed with PV- or SOM-specific oIPSCs. Cd decreased amplitude in both cases (Norm oIPSC; PV = 0.29 ± 0.05, n/N = 6/2; p = 0.31; SOM = 0.36 ± 0.07, n/N = 7/2; p = 0.016, Wilcoxon signed rank test) without increasing PPR (Norm PPR; PV = 0.98 ± 0.06; p = 0.84; SOM = 0.97 ± 0.05; p = 0.69, Wilcoxon signed rank test; **Extended Data 2E-F**). This indicates that both PV+ and SOM+ terminals exhibit nanodomain configurations, opening the possibility that DOR engages non-canonical gain modulation in SOM+ terminals.

### PV+ interneurons exhibit canonical presynaptic Ca_V_ inhibition

Before assessing the complexities of SOM+ terminals, we first wanted to validate that DORs engage exclusively canonical signaling cascades in PV+ terminals. DORs, like other opioid receptors, are classically coupled to inhibitory G_i/o_ -proteins^21,48^ that inhibit presynaptic Ca_V_ s directly via translocated Gβγ subunits^22,25,49-51^. Since DOR modulation of PV-oIPSCs overlapped with the low Ca relationship (**Figure 2E-F**), Gβγ-dependent Ca_V_ inhibition presumably underlies the decrease in *P*_*R*_ ^27,28,48^, as observed in hippocampal basket cells^37^. However, a recent study found that mu opioid receptors (MORs) suppress PV-oIPSCs in orbitofrontal cortex through cAMP-dependent protein kinase (PKA)^31^. To test this in mPFC, we recorded PV-oIPSCs with the selective PKA inhibitor H89 (10 μM) included in the ACSF (**Figure 3A**). Following DPDPE, oIPSC depression was comparable to control (i.e., DPDPE alone) (Norm Amp; control vs. H89 = 0.34 ± 0.04 vs. 0.45 ± 0.06; p = 0.13, unpaired t-test; **Figure 3B**) as was the amplitude-PPR relationship (control vs. H89 slope = -0.50 ± 0.07 vs. -0.66 ± 0.16; p = 0.73, Dunnett’s test; **Figure 3C**). H89 did not affect PV-oIPSC amplitudes at baseline (control vs. H89 = -800.9 ± 116.2 pA vs. -680.6 ± 122.8 pA, n/N = 12/5 vs. 8/3; p = 0.49, unpaired t-test), indicating that PKA did not regulate basal GABA release. This demonstrated that presynaptic DOR signaling in PV+ neurons does not require PKA, and instead is likely mediated by Gβγ-mediated mechanisms.

**Figure 3:**
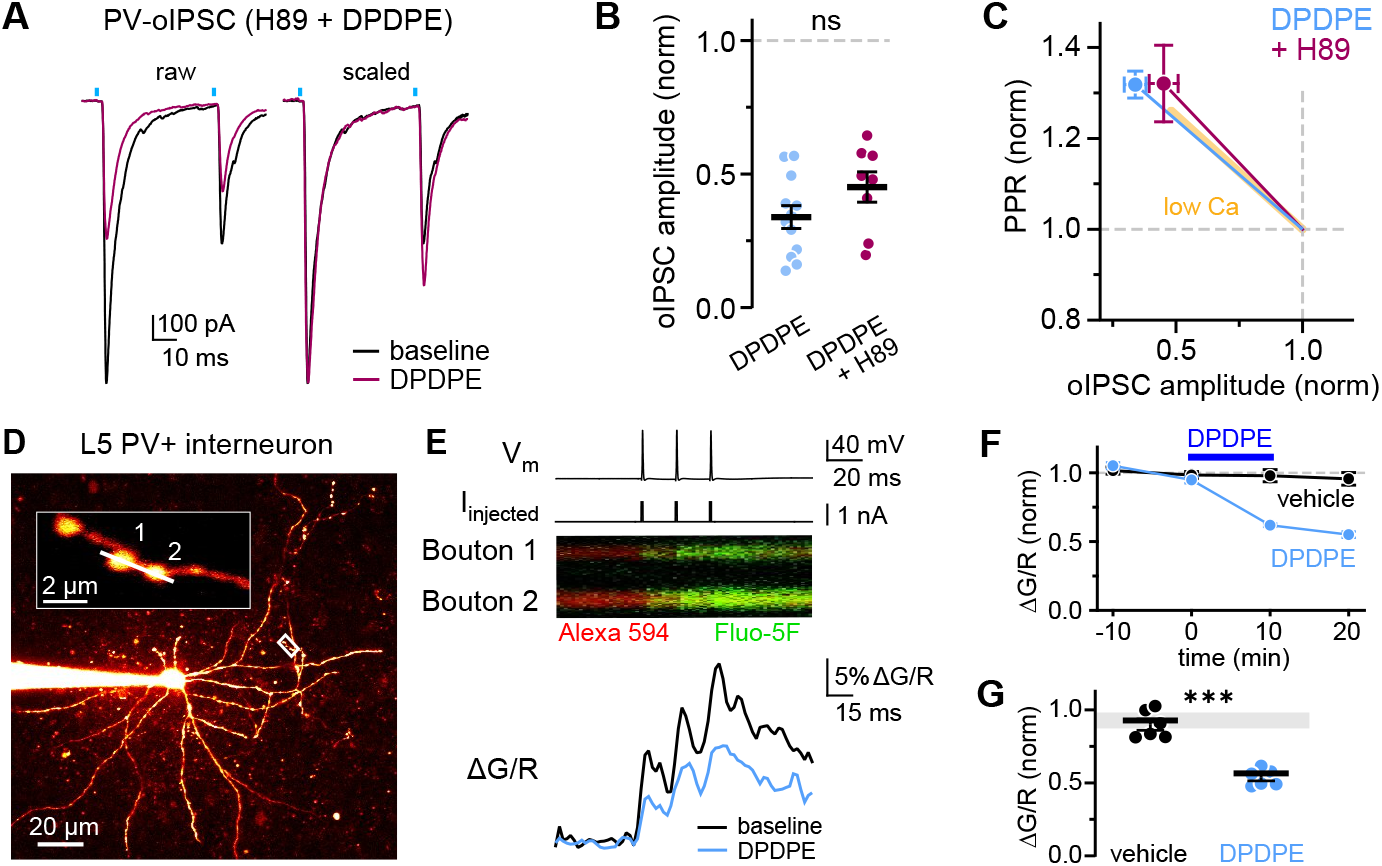
DORs mediate canonical Ca_V_ inhibition in PV+ interneurons. **A:** Example PV-oIPSC traces before and after application of DPDPE while in the presence of H89 (10 μM). **B:** Summary plot of normalized PV-oIPSC amplitudes. **C:** Summary amplitude-PPR plot depicting slope relationships of PV-oIPSCs after DPDPE. Low Ca eIPSC relationship (yellow line) from Figure 1 is included for comparison. **D:** Z-stack image of filled L5 PV+ interneuron. White box indicates bouton imaging area, which is enlarged and rotated as a zoomed-in inset (top left). Linescan ROI is represented as white bar. Scale bars, main image: 20 μm, inset: 2 μm. **E:** Top: Raw voltage trace of AP triplet evoked by step protocol (middle). Middle: time vs. fluorescence intensity image of superimposed Alexa 594 (red) and Fluo-5F (green) signals from both boutons. Bottom: Example Ca transients from ΔG/R fluorescence data at baseline and following DPDPE application. **F:** Summary plot of normalized ΔG/R peaks over time averaged over multiple PV+ cells. **G:** Summary plot of normalized ΔG/R peaks in vehicle and DPDPE conditions. Normalized ΔG/R peaks are the final timepoint (20 min after DPDPE application) relative to baseline average. *** p<0.001.

If DORs suppress release by altering presynaptic Ca_V_ function, this should be evident by imaging Ca influx at terminals. To test this, mPFC slices were obtained from fluorescent reporter mice (PV-Ai14) that allowed identification of PV+ somata via 2-photon microscopy. Whole-cell patch clamp recordings were made from fluorescent cells in L5, and cells were loaded with the volume-filling dye AlexaFluor 594 (20 μM) and the Ca indicator Fluo-5F (250 μM) (**Figure 3D**). PV+ interneuron identity was validated through neurophysiological characterization (low R_input_, high-frequency AP firing with pronounced AP after-hyperpolarization). A burst of 3 APs was then evoked via step current injection (3x at 50 Hz, 1-2 nA, 1 ms) and resultant AP-evoked Ca influx was imaged at boutons (**Figure 3E**). Robust Ca transients were elicited reliably by this protocol and were stable over 40 minutes in control conditions. Following application of DPDPE, AP-evoked Ca transients decreased significantly (Norm ΔG/R; vehicle vs. DPDPE = 0.90 ± 0.04 vs. 0.54 ± 0.03, n/N = 6/3 vs. 6/3; p = 0.00011, unpaired t-test; **Figure 3F-G**), demonstrating that DORs inhibit presynaptic Ca_V_ in PV+ terminals. Additionally, DPDPE lowered R_input_ (ΔR_input_; vehicle vs. DPDPE = 0.9 ± 2.4 % vs. -5.3 ± 2.2 %, n/N = 6/3 vs. 6/3; p = 0.028, unpaired t-test; **Extended Data 3B**) and increased outward holding current (ΔI_hold_; vehicle vs. DPDPE = 0.5 ± 3.8 pA vs. 30.5 ± 5.6 pA; p = 0.0004, unpaired t-test; **Extended Data 3C**) corresponding to membrane hyperpolarization, likely through GIRK activation^37^. These results demonstrate that DORs strongly inhibit PV+ output, influencing both somatodendritic and axonal compartments, and that DOR suppresses release from PV+ terminals by modulating Ca_V_s.

### DORs engage multiple modulatory cascades in SOM+ neurons

Excitatory synapses in mPFC express two forms of presynaptic modulation: a canonical G_i/o_-Gβγ pathway (i.e., GABA_B_R) and non-canonical signaling via G_s_ and PKA (i.e., D1R)^35^. DOR-dependent modulation in SOM+ terminals, by contrast, appear intermediate between these two mechanisms. Thus, we hypothesize that DORs engage two independent signaling cascades in SOM+ terminals that converge on Ca_V_s. If true, then each signaling cascade should contribute partially to the total reduction in IPSC amplitude, but with different effects on PPR. These different effects could then sum to yield an intermediate PPR result that lies between canonical and gain modulation (**Figure 2C**).

Testing this requires selective block of the two putative signalling cascades driven by G_i/o_-Gβγ and G_s_-PKA. While PKA activity can be easily blocked in slice preparations, blockade of G_i/o_-Gβγ is more difficult. Pertussis toxin (PTx)—a highly potent and specific inhibitor of G_i/o_-coupled signaling—requires prolonged exposure (>24 hrs) to achieve a saturating effect in tissue^52,53^, making it unwieldy for traditional slice experiments. To achieve this saturating block, we delivered PTx via stereotaxic injection locally to mPFC, 1-3 days prior to slice preparation, in SOM-Cre mice previously infected with ChR2-expressing viruses (**Figure 4A**). This strategy was chosen over other delivery methods due to its increased efficacy (see Methods; but see also^49,50^). To confirm that G_i/o_ -Gβγ signaling was attenuated by this method, we assayed GABA_B_-dependent modulation of electrically-evoked EPSCs with 1 μM baclofen in one slice from each animal. Baclofen reduced eEPSCs by 77% in uninjected slices (n/N = 5/2, p < 0.0001, paired t-test), comparable to prior observations^23,54^. By contrast, baclofen reduced eEPSCs by only 10% in slices from PTx treated animals (Norm EPSC: 0.90 ± 0.02, n/N = 10/8; p < 0.0001, unpaired t-test; **Figure 4B**), indicating that the majority of G_i/o_ signaling was blocked with this approach.

**Figure 4:**
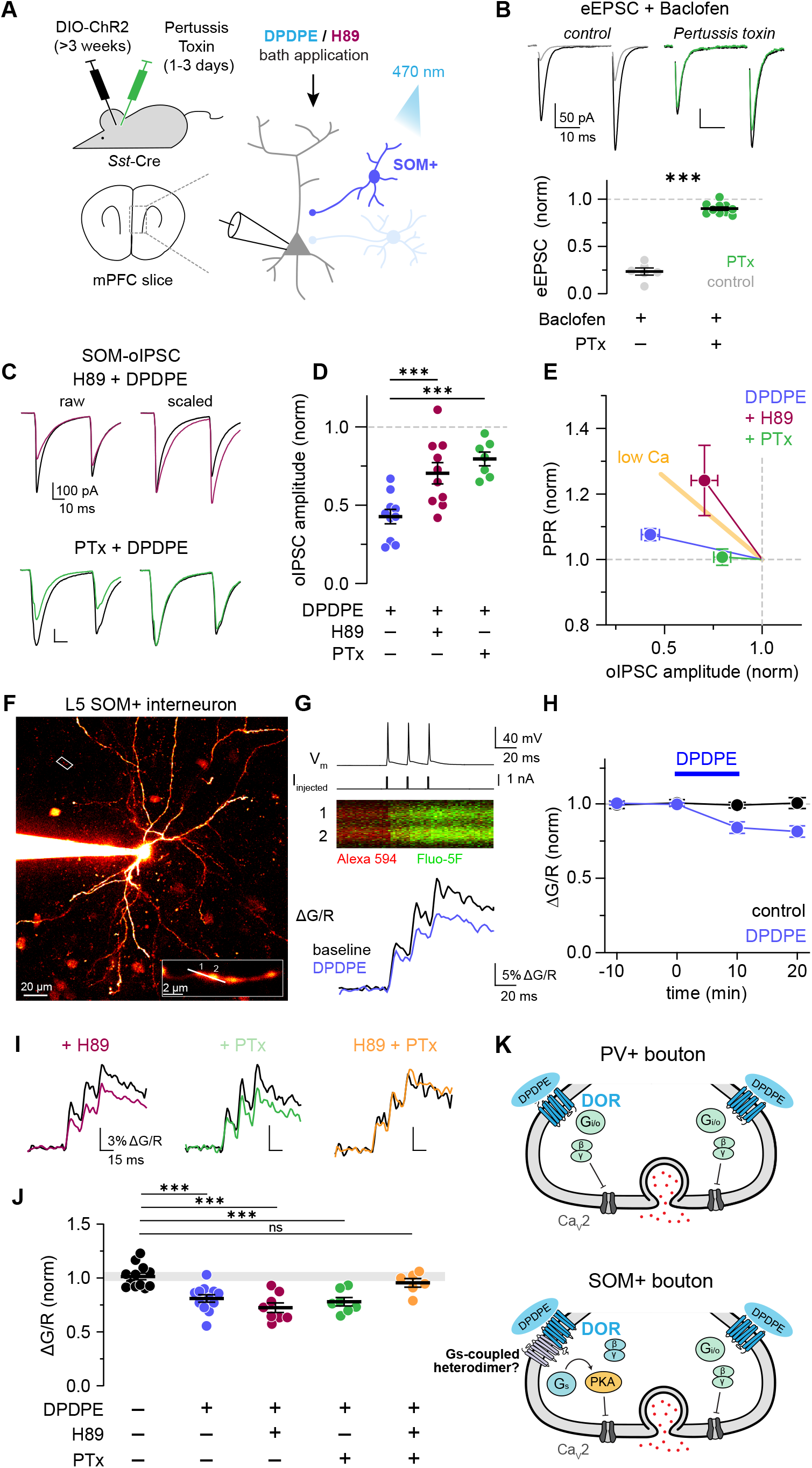
SOM+ interneurons exhibit heterogeneous regulation by DORs. **A:** Cartoon depicting injection schedule for PTx electrophysiology experiments. Two stereotaxic surgeries are performed on SOM-Cre mice; slices are obtained 3-4 weeks after injection of DIO-ChR2 and 1-3 days after injection of PTx (0.5 μg/ μL). **B:** Example EPSC traces in L5 pyramidal neurons evoked by electrical stimulation before and after application of baclofen (1 μM) in control mice (left) or those locally-injected with PTx (right). Below: Summary plot of normalized eEPSC amplitudes after baclofen application. **C:** Example SOM-oIPSC traces before and after DPDPE with either external H89 (top) or injected PTx (bottom). **D:** Summary plot of normalized oIPSC amplitudes after DPDPE in control, H89, and PTx conditions. **E:** Normalized amplitude-PPR plot depicting slope relationships for control SOM-oIPSCs after DPDPE, compared to those in the presence of H89 or PTx. Yellow line depicts benchmark low Ca eIPSC relationship from Figure 1. **F:** Z-stack image of filled SOM+ interneuron. White box indicates the bouton imaging region, which is enlarged and rotated as a zoomed-in inset (bottom). Scale bars, main image: 20 μm, inset: 2 μm. **G:** Top: Voltage trace of AP triplet evoked by step protocol (middle). Bottom: time vs. fluorescence intensity image of Alexa 594 (red) and Fluo-5F (green) signals. **H:** Summary plot of normalized ΔG/R peaks over time comparing vehicle and DPDPE group averages. **I:** Example AP-evoked Ca transients before and after DPDPE with either H89, locally-injected PTx, or both. **J:** Summary plot of normalized ΔG/R peaks across vehicle and drug conditions. *** p<0.001. **K:** Cartoon depicting DOR-dependent signaling pathway in PV+ and SOM+ boutons.

Following PTx validation, SOM-oIPSCs were evoked as described earlier (**Figure 2**). In these conditions, DPDPE application still reduced oIPSCs in PTx-injected slices (Norm oIPSC = 0.80 ± 0.04, n/N = 7/3; p = 0.0034, paired t-test), though to a lesser degree compared to DPDPE application in the absence of PTx (p = 0.0006, Holm-Sidak post-hoc test; **Figure 4C-D**). Similar effects were observed with the PKA blocker, H89 (Norm oIPSC; control vs. H89 = 0.43 ± 0.05 vs. 0.70 ± 0.07, n/N = 10/4 vs. 10/4; p = 0.0025, Holm-Sidak post-hoc test; **Figure 4C-D**). Strikingly, the amplitude-PPR relationships were completely distinct for PTx and H89 conditions. With PKA blocked, the residual modulation was completely canonical, overlaying with low Ca. With G_i/o_-Gβγ blocked, PPR was not altered (**Figure 4E**). Taken together, these results demonstrate that DORs regulate SOM+ GABA release through two mechanisms: one that is G_i/o_-dependent that increases PPR, and another that is G_i/o_-independent but PKA-dependent, that does not alter PPR.

While the data above indicate that DORs engage two separate signaling cascades to regulate release from SOM+ terminals. But since IPSCs are the aggregate of activity across multiple synapses, they cannot inform whether both cascades are engaged within single terminals or are engaged selectively on a terminal-by-terminal basis, perhaps based on SOM+ cell subclass. Somatostatin is expressed by a range of interneuron subclasses that can be grouped by intrinsic electrophysiological characteristics^55-57^. Indeed, when recording directly from SOM+ cells, we identified two distinct neurophysiological phenotypes, with some neurons exhibiting high R_input_, slow τ_membrane_, and small afterhyperpolarizations (AHP), and others with lower R_input_, faster τ_membrane_, larger AHP, and quasi-fast-spiking behavior akin to PV+ neurons (**Extended Data 4B**). These two subgroups are defined as putative Martinotti (MC) and non-Martinotti (NMC) cells^58^, and contrast with PV+ neurons further by repetitive spike patterns and AP waveform (**Extended Data 4D**). We therefore made whole-cell recordings from each of these classes and imaged AP-evoked Ca influx in boutons (**Figure 4F-G**). Ca transients were reduced by DPDPE in all boutons imaged, independent of SOM subclass (Norm ΔG/R in MC; vehicle vs. DPDPE = 1.00 ± 0.03 vs. 0.81 ± 0.03, n/N = 11/9 vs. 12/8; p = 0.0007, Holm-Sidak post-hoc test; **Figure 4H, Extended Data 4G**). Remarkably, DPDPE was similarly effective in all boutons imaged in the presence of H89 or PTx (Norm ΔG/R; control vs. H89 = 0.81 ± 0.03 vs. 0.73 ± 0.05, n/N = 11/9 vs. 8/5; p = 0.36; control vs. PTx = 0.81 ± 0.03 vs. 0.78 ± 0.04, n/N = 11/9 vs. 7/5; p = 0.66, Holm-Sidak post-hoc test; **Figure 4I-J**). Combined application of both H89 and PTx completely eliminated the effect of DPDPE on Ca transients in all SOM+ boutons imaged (Norm ΔG/R; vehicle vs. H89 + PTx = 1.00 ± 0.03 vs. 0.96 ± 0.04, n/N = 11/9 vs. 6/4; p = 0.66, Holm-Sidak post-hoc test; **Figure 4I-J**). Taken together, these results indicate that DORs engage two distinct DOR signaling pathways within all SOM+ boutons, independent of SOM subclass, and that both cascades converge on presynaptic Ca_V_.

## DISCUSSION

Here we find that DORs suppress prefrontal GABA release through multiple mechanisms that are specific to inhibitory neuron subclass. While canonical G_i/o_-Gβγ signaling predominates in PV+ presynaptic terminals, SOM+ boutons unexpectedly exhibit heterogeneous DOR signaling, with both G_i/o_-dependent and -independent pathways present at individual presynaptic terminals. Both ultimately inhibit Ca influx through presynaptic Ca_V_s. G_i/o_-Gβγ signaling depolarizes the voltage-dependence of channel activation. G_i/o_-Gβγ-independent, PKA-dependent signaling, as shown previously^35^, instead reduces the probability that channels will open at all in response to an AP. We found here that these two mechanisms likely act in parallel, in the same boutons, presumably because these mechanisms can multiplex to affect different aspects of presynaptic Ca_V_ function.

PV+ and SOM+ inputs target different compartments of individual L5 pyramidal neurons^59^. Since STP dictates the frequency-dependence of transmission, each afferent type is subject to different levels of temporal filtering by DORs. G_i/o_-Gβγ modulation regulates PV+ inputs that target the soma and axon initial segment (AIS). Canonical STP changes caused by DOR activation therefore impose a high-pass filter of GABAergic transmission at these inputs, with strong suppression of APs that occur in isolation and facilitation of events that occur at high-frequency. This means that highly active PV+ inputs can overcome DOR-induced suppression and the level of inhibitory tone on perisomatic compartments may remain relatively stable, promoting consistent inhibition of AP generation^60^. By contrast, SOM+ inputs primarily target more distal dendritic regions and exert strong control over local signal integration^61^. DORs affect STP to a lesser extent at these inputs, making SOM-mediated inhibition particularly resistant to frequency-dependent temporal filtering. Therefore, DORs at SOM+ terminals can promote a more consistent disinhibition of distal pyramidal cell dendritic segments, even during periods of high SOM+ cell activity, relative to perisomatic compartments.

### Pleiotropic opioid receptor signaling

Presynaptic opioid receptors engage a variety of intracellular cascades throughout the brain. Although G_i/o_ (and specifically G_i/o_-Gβγ) signaling is common, other pathways mobilize β-arrestin or G_q_-subunits that may regulate HCN or K_V_ channels instead of Ca_V_ and GIRK^27,62^. Despite this diversity, one may assume that an opioid receptor agonist at a given synapse will engage a single signaling mode that yields a consistent, predictable effect on synaptic transmission. Data shown here and at other synapses^63^ suggest that this may be an oversimplification. For example, after chronic morphine treatment, DORs in the nucleus raphe magnus inhibit GABA release through both phospholipase A2 and cAMP/PKA signaling acting in parallel^64^. In the ventral tegmental area, acute morphine application reduces IPSC frequency through MORs via β-arrestin and c-Src tyrosine kinase^65^, while MOR-dependent G_i/o_ /cAMP signaling has also been shown to suppress GABA release at these synapses^66^.

Pleiotropic signaling that regulates transmitter release via simultaneous, parallel pathways may be more common than previously appreciated. If opioid receptors regulate transmitter release by reducing *P*_*R*_, PPR should consistently and proportionately increase during opioid-mediated suppression. This assumption does not always hold true. Multiple studies of opioid-dependent synaptic regulation suggest that modulation occurs presynaptically, even when PPR changes only slightly or not at all^31-34,67^. While there are Ca_V_ -independent mechanisms that could account for some aspects of these effects^68,69^, we show here that parallel signaling cascades converging on Ca_V_s can account for modulation entirely in these GABAergic terminals (**Figure 4**).

Revisiting prior findings through this lens suggests that this pleiotropic signaling may be engaged at multiple synapses by multiple opioid receptors. Studies often conclude that GPCRs are acting presynaptically if *any* increase in PPR is observed. But here we show that the magnitude of PPR change, benchmarked against how much it should change via canonical mechanisms, can inform on *how* Ca_V_s are being modulated. As a case in point, DORs at nucleus raphe magnus synapses reduces EPSC amplitude ∼50% while PPR increases by only 12%^64^, a change far too small to be explained via canonical mechanisms. There are comparable observations found elsewhere, including CA1^37,70^, subthalamic nucleus^38^, and intercalated cells of the amygdala^71^, where GPCRs cause substantial synaptic depression with only modest, yet statistically significant, effects on PPR. That intermediate PPR change can result from the combination of multiple pathways has implications for interpreting presynaptic opioid effects more broadly and suggests that a new framework is necessary.

### Presynaptic gain modulation

We previously demonstrated that transmitter release can be linearly tuned via presynaptic gain modulation^35^. D1R gain modulation shares hallmark features with G_i/o_-independent regulation by DORs observed here, including stable STP, PKA dependency, and Ca_V_ dependency. This paradigm provides the best explanation for noncanonical DOR signaling in SOM+ boutons and could account for other cases where opioid receptors appear to act atypically^31-34,67^. These mechanisms add to our understanding of non-canonical pathways for regulation of transmission that, in the past, have focused on Ca_V_-independent components. For example, following receptor activation, dissociated Gβγ can bind directly to SNAP25, a component of the presynaptic SNARE complex, which prevents membrane fusion and vesicle exocytosis independent of Ca influx^72^. G_i/o_ -coupled 5-HT_1B_ receptors depress synaptic release via Gβγ-SNAP25 interactions in the hippocampus^73^ and spinal cord^68^. Individual terminals express multiple types of G_i/o_-coupled receptors that engage either Gβγ-SNAP25 or canonical Ca_V_ regulation, as in the hippocampus^73^ and nucleus accumbens^74^. For example, MORs suppress excitatory transmission in nucleus accumbens solely via Gβγ-SNAP25 interaction, while kappa receptors (KORs) utilize a different pathway to regulate the same inputs^74^. In the bed nucleus of the stria terminalis, KORs suppress GABA release from amygdala inputs without causing an increase in STP^32^. Instead of Gβγ-SNAP25 or PKA-Ca_V_, KOR modulation required extracellular signal-regulated kinase (ERK), distinguishing it from the types of signaling previously discussed. Taken together, presynaptic regulation of transmitter release can be achieved through multiple opioid receptor pathways, with different mechanisms invoking different temporal filtering characteristics on transmission.

### G_i/o_-independent opioid receptor signaling

In SOM+ boutons, we observed a signaling component downstream of DOR activation that persisted in the presence of PTx. Since opioid receptors are most often coupled with PTx-sensitive G_i/o_ subunits, DORs must be engaging a separate pathway. Opioid receptors can interact with the atypical G_i/o_ family subunit G_z_, which, while still reducing levels of cAMP^21^, is notably insensitive to PTx^75^. G_z_ subunits interact with DORs *in vitro*^76^, but they may preferentially associate with MORs in the brain^77^ and have not been found presynaptically, making their presence in SOM+ boutons unlikely. Furthermore, we demonstrated that PKA activation is required for gain-style DOR regulation since H89 blocks this component entirely (**Figure 4**). If DORs were acting through G_i/o_ signaling, application of H89 should have enhanced the IPSC depression caused by DPDPE since both lead to PKA inhibition. Instead, the gain component of DOR modulation is most likely mediated via G_s_ signaling (PKA-dependent, PTx-insensitive), as with D1R gain modulation^35^.

How could G_i/o_ -coupled DORs signal via G_s_ ^78,79^? The most likely mechanism is that DORs heterodimerize with G_s_-coupled receptors in SOM+ terminals. Heterodimeric GPCR complexes consist of two different receptor protomers that oligomerize either directly or through a shared membrane effector (i.e., adenylyl cyclase (AC))^80^. This broadens the regulatory capacity of a receptor complex and confers signaling properties distinct from individual subunits. DORs can form heterodimers and have been shown to oligomerize with both MORs and KORs^81,82^, as well as with cannabinoid and chemokine receptors^83,84^, affecting desensitization, pharmacological efficacy, and signaling bias. G_s_ and G_i/o_ receptors, though having opposed downstream effects, can also oligomerize through shared AC proteins, yielding distinctive signaling properties. For example, G_s_-coupled adenosine A_2A_ and G_i/o_-coupled dopamine D2 receptors form a common functional unit with AC5 proteins, each exerting allosteric control over catalyzation of cAMP^85^. DORs have also been shown to co-immunoprecipitate with G_s_-coupled β2 adrenergic receptors in cardiac tissue^86^, establishing physiological precedent for DOR-containing G_s_ -G_i/o_ heterodimers.

GPCR heterodimers are attractive therapeutic targets, potentially offering tissue-specificity and limited side effects compared to receptor protomers or homodimers^80^. Selective pharmacology has been developed for a small subset of discovered heterodimer pairs, including mGluR_2/4_ receptors^87^. A combination of positive and negative allosteric modulators was used to show that mGluR heteromers, but not homomers, regulate thalamic inputs to mPFC, but not those from hippocampus or amygdala^88^. Targeting peripheral MOR/DOR or MOR/KOR heterodimers has shown promise for producing analgesia with less tolerance and withdrawal effects, but the functional relevance of opioid receptor heterodimers in the brain is less well-characterized^89^. Thus, efforts should be made in the future to both determine whether DORs engage PKA through a heterodimer complex, determine the identify of this putative G_s_-coupled partner, and explore pharmacological methods to selectively regulate its activity independent of DOR protomers.

### Functional implications

DORs evoke differential, pathway-specific modulation of inhibitory inputs to pyramidal cells. Ultimately, DORs suppress GABA release from both interneuron subclasses and would thus broadly have disinhibitory influence. However, since DORs are also expressed at glutamatergic synapses in PFC^39^, the net effect on excitation-inhibition balance is harder to predict and may depend on afferent source, as is the case for KOR signaling^90^. Both PV+ and SOM+ neurons exhibit strong presynaptic regulation by DORs, which would have compartment-specific effects in individual pyramidal cells. Perisomatic-targeting PV+ neurons are important for establishing synchronous firing, particularly in the gamma range (30-80 Hz)^91^, of pyramidal cell networks in the neocortex and hippocampus^92^. Presynaptic DOR regulation at these synapses—mediated through canonical G_i/o_-Gβγ signaling—imposes a high-pass filter on GABA release; therefore, the strength of inhibitory drive onto perisomatic compartments will recover during longer and higher frequency stimuli. Brief disinhibition via DORs could tune the oscillatory phase of pyramidal activity on shorter timescales, as very few PV+ APs can control spike-timing^93^, while preventing runaway network excitation on longer timescales^94^. Comparatively, dendrite-targeting SOM+ neurons have less direct control over AP initiation and instead modulate local dendritic excitability. In L5 neurons, this may affect generation and propagation of dendritic Ca spikes, the large amplitude distal depolarizations associated with somatic burst firing thought to be important for integrating signals from multiple brain areas^95^. SOM-derived GABA has been shown to suppress dendritic spikes^96^, so local disinhibition by DORs may increase the likelihood of such events and promote associative plasticity^97^. Such disinhibition and expansion of plasticity associative timing windows has been observed for D2 dopaminergic regulation of interneurons in prefrontal cortex^98^. DOR-based regulation, with its bias towards disinhibiting SOM+ inputs, could serve as a mechanism to promote associative plasticity selectively of long-range synaptic inputs that converge on apical dendrites. Given that DOR-mediated disinhibition at SOM+ terminals is less affected by presynaptic firing frequency, plasticity could be facilitated even during periods of elevated circuit activity. New intersectional genetic tools (i.e., conditional DOR deletion in either PV+ or SOM+ neurons) will be required to fully elucidate the role of differential DOR disinhibition on pyramidal cell computation and prefrontal circuit processing.

## Supplementary materials

Supplementary materials contain Methods and Extended Figures 1-5.

## Acknowledgments

We are grateful to members of the Bender lab for comments and feedback. We thank Dr. Selin Schamiloglu for assistance with imaging analysis, Dr. Guy Bouvier for initial access to SOM-Cre mice, Dr. Elyssa Margolis for discussions regarding DOR pharmacology, and Drs. Dorit Ron and Jennifer Whistler for discussions related to G_s_-G_i/o_ heterodimers. This work was supported by NIH Grants MH112729 and AA027023 (K.J.B.).

## Author contributions

R.P.D.A. and K.J.B. conceived the project and designed the experiments. R.P.D.A. performed all experiments and data analysis. R.P.D.A. and K.J.B. wrote the manuscript.

## REFERENCES

1. Mansour, A., Fox, C.A., Akil, H., and Watson, S.J. (1995). Opioid-receptor mRNA expression in the rat CNS: anatomical and functional implications. Trends Neurosci 18, 22–29. 10.1016/0166-2236(95)93946-u.

2. Waldhoer, M., Bartlett, S.E., and Whistler, J.L. (2004). Opioid receptors. Annu Rev Biochem 73, 953–990. 10.1146/annurev.biochem.73.011303.073940.

3. Le Merrer, J., Becker, J.A., Befort, K., and Kieffer, B.L. (2009). Reward processing by the opioid system in the brain. Physiol Rev 89, 1379–1412. 10.1152/physrev.00005.2009.

4. Contet, C., Kieffer, B.L., and Befort, K. (2004). Mu opioid receptor: a gateway to drug addiction. Curr Opin Neurobiol 14, 370–378. 10.1016/j.conb.2004.05.005.

5. Kieffer, B.L., and Gaveriaux-Ruff, C. (2002). Exploring the opioid system by gene knockout. Prog Neurobiol 66, 285–306. 10.1016/s0301-0082(02)00008-4.

6. Chu Sin Chung, P., and Kieffer, B.L. (2013). Delta opioid receptors in brain function and diseases. Pharmacol Ther 140, 112–120. 10.1016/j.pharmthera.2013.06.003.

7. Pradhan, A.A., Befort, K., Nozaki, C., Gaveriaux-Ruff, C., and Kieffer, B.L. (2011). The delta opioid receptor: an evolving target for the treatment of brain disorders. Trends Pharmacol Sci 32, 581–590. 10.1016/j.tips.2011.06.008.

8. Quirion, B., Bergeron, F., Blais, V., and Gendron, L. (2020). The Delta-Opioid Receptor; a Target for the Treatment of Pain. Front Mol Neurosci 13, 52. 10.3389/fnmol.2020.00052.

9. Margolis, E.B., Fields, H.L., Hjelmstad, G.O., and Mitchell, J.M. (2008). Delta-opioid receptor expression in the ventral tegmental area protects against elevated alcohol consumption. J Neurosci 28, 12672–12681. 10.1523/JNEUROSCI.4569-08.2008.

10. Nielsen, C.K., Simms, J.A., Pierson, H.B., Li, R., Saini, S.K., Ananthan, S., and Bartlett, S.E. (2008). A novel delta opioid receptor antagonist, SoRI-9409, produces a selective and long-lasting decrease in ethanol consumption in heavy-drinking rats. Biol Psychiatry 64, 974–981. 10.1016/j.biopsych.2008.07.018.

11. Chefer, V.I., and Shippenberg, T.S. (2009). Augmentation of morphine-induced sensitization but reduction in morphine tolerance and reward in delta-opioid receptor knockout mice. Neuropsychopharmacology 34, 887–898. 10.1038/npp.2008.128.

12. Scherrer, G., Imamachi, N., Cao, Y.Q., Contet, C., Mennicken, F., O’Donnell, D., Kieffer, B.L., and Basbaum, A.I. (2009). Dissociation of the opioid receptor mechanisms that control mechanical and heat pain. Cell 137, 1148–1159. 10.1016/j.cell.2009.04.019.

13. Chu Sin Chung, P., Keyworth, H.L., Martin-Garcia, E., Charbogne, P., Darcq, E., Bailey, A., Filliol, D., Matifas, A., Scherrer, G., Ouagazzal, A.M., et al. (2015). A novel anxiogenic role for the delta opioid receptor expressed in GABAergic forebrain neurons. Biol Psychiatry 77, 404–415. 10.1016/j.biopsych.2014.07.033.

14. Chung, P.C., Boehrer, A., Stephan, A., Matifas, A., Scherrer, G., Darcq, E., Befort, K., and Kieffer, B.L. (2015). Delta opioid receptors expressed in forebrain GABAergic neurons are responsible for SNC80-induced seizures. Behav Brain Res 278, 429–434. 10.1016/j.bbr.2014.10.029.

15. Saitoh, A., Suzuki, S., Soda, A., Ohashi, M., Yamada, M., Oka, J.I., Nagase, H., and Yamada, M. (2018). The delta opioid receptor agonist KNT-127 in the prelimbic medial prefrontal cortex attenuates veratrine-induced anxiety-like behaviors in mice. Behav Brain Res 336, 77–84. 10.1016/j.bbr.2017.08.041.

16. Baldo, B.A. (2016). Prefrontal Cortical Opioids and Dysregulated Motivation: A Network Hypothesis. Trends Neurosci 39, 366–377. 10.1016/j.tins.2016.03.004.

17. Koob, G.F. (2020). Neurobiology of Opioid Addiction: Opponent Process, Hyperkatifeia, and Negative Reinforcement. Biol Psychiatry 87, 44–53. 10.1016/j.biopsych.2019.05.023.

18. Jiang, C., Wang, X., Le, Q., Liu, P., Liu, C., Wang, Z., He, G., Zheng, P., Wang, F., and Ma, L. (2021). Morphine coordinates SST and PV interneurons in the prelimbic cortex to disinhibit pyramidal neurons and enhance reward. Mol Psychiatry 26, 1178–1193. 10.1038/s41380-019-0480-7.

19. Olive, M.F., Anton, B., Micevych, P., Evans, C.J., and Maidment, N.T. (1997). Presynaptic versus postsynaptic localization of mu and delta opioid receptors in dorsal and ventral striatopallidal pathways. J Neurosci 17, 7471–7479. 10.1523/JNEUROSCI.17-19-07471.1997.

20. Ostermeier, A.M., Schlosser, B., Schwender, D., and Sutor, B. (2000). Activation of mu- and delta-opioid receptors causes presynaptic inhibition of glutamatergic excitation in neocortical neurons. Anesthesiology 93, 1053–1063. 10.1097/00000542-200010000-00029.

21. Che, T., and Roth, B.L. (2023). Molecular basis of opioid receptor signaling. Cell 186, 5203–5219. 10.1016/j.cell.2023.10.029.

22. Morikawa, H., Mima, H., Uga, H., Shoda, T., and Fukuda, K. (1999). Opioid potentiation of N-type Ca2+ channel currents via pertussis-toxin-sensitive G proteins in NG108-15 cells. Pflugers Arch 438, 423–426. 10.1007/pl00008091.

23. Mintz, I.M., and Bean, B.P. (1993). GABAB receptor inhibition of P-type Ca2+ channels in central neurons. Neuron 10, 889–898. 10.1016/0896-6273(93)90204-5.

24. Bean, B.P. (1989). Neurotransmitter inhibition of neuronal calcium currents by changes in channel voltage dependence. Nature 340, 153–156. 10.1038/340153a0.

25. Herlitze, S., Garcia, D.E., Mackie, K., Hille, B., Scheuer, T., and Catterall, W.A. (1996). Modulation of Ca2+ channels by G-protein beta gamma subunits. Nature 380, 258–262. 10.1038/380258a0.

26. Al-Hasani, R., and Bruchas, M.R. (2011). Molecular mechanisms of opioid receptor-dependent signaling and behavior. Anesthesiology 115, 1363–1381. 10.1097/ALN.0b013e318238bba6.

27. Reeves, K.C., Shah, N., Munoz, B., and Atwood, B.K. (2022). Opioid Receptor-Mediated Regulation of Neurotransmission in the Brain. Front Mol Neurosci 15, 919773. 10.3389/fnmol.2022.919773.

28. Atwood, B.K., Lovinger, D.M., and Mathur, B.N. (2014). Presynaptic long-term depression mediated by Gi/o-coupled receptors. Trends Neurosci 37, 663–673. 10.1016/j.tins.2014.07.010.

29. Lovinger, D.M., Mateo, Y., Johnson, K.A., Engi, S.A., Antonazzo, M., and Cheer, J.F. (2022). Local modulation by presynaptic receptors controls neuronal communication and behaviour. Nat Rev Neurosci 23, 191–203. 10.1038/s41583-022-00561-0.

30. Jackman, S.L., and Regehr, W.G. (2017). The Mechanisms and Functions of Synaptic Facilitation. Neuron 94, 447–464. 10.1016/j.neuron.2017.02.047.

31. Lau, B.K., Ambrose, B.P., Thomas, C.S., Qiao, M., and Borgland, S.L. (2020). Mu-Opioids Suppress GABAergic Synaptic Transmission onto Orbitofrontal Cortex Pyramidal Neurons with Subregional Selectivity. J Neurosci 40, 5894–5907. 10.1523/JNEUROSCI.2049-19.2020.

32. Li, C., Pleil, K.E., Stamatakis, A.M., Busan, S., Vong, L., Lowell, B.B., Stuber, G.D., and Kash, T.L. (2012). Presynaptic inhibition of gamma-aminobutyric acid release in the bed nucleus of the stria terminalis by kappa opioid receptor signaling. Biol Psychiatry 71, 725–732. 10.1016/j.biopsych.2011.11.015.

33. Tejeda, H.A., Wu, J., Kornspun, A.R., Pignatelli, M., Kashtelyan, V., Krashes, M.J., Lowell, B.B., Carlezon, W.A., Jr., and Bonci, A. (2017). Pathway- and Cell-Specific Kappa-Opioid Receptor Modulation of Excitation-Inhibition Balance Differentially Gates D1 and D2 Accumbens Neuron Activity. Neuron 93, 147–163. 10.1016/j.neuron.2016.12.005.

34. Yokota, E., Koyanagi, Y., Yamamoto, K., Oi, Y., Koshikawa, N., and Kobayashi, M. (2016). Opioid subtype- and cell-type-dependent regulation of inhibitory synaptic transmission in the rat insular cortex. Neuroscience 339, 478–490. 10.1016/j.neuroscience.2016.10.004.

35. Burke, K.J., Jr., Keeshen, C.M., and Bender, K.J. (2018). Two Forms of Synaptic Depression Produced by Differential Neuromodulation of Presynaptic Calcium Channels. Neuron 99, 969–984 e967. 10.1016/j.neuron.2018.07.030.

36. van Rijn, R.M., Defriel, J.N., and Whistler, J.L. (2013). Pharmacological traits of delta opioid receptors: pitfalls or opportunities? Psychopharmacology (Berl) 228, 1–18. 10.1007/s00213-013-3129-2.

37. He, X.J., Patel, J., Weiss, C.E., Ma, X., Bloodgood, B.L., and Banghart, M.R. (2021). Convergent, functionally independent signaling by mu and delta opioid receptors in hippocampal parvalbumin interneurons. Elife 10. 10.7554/eLife.69746.

38. Shen, K.Z., and Johnson, S.W. (2002). Presynaptic modulation of synaptic transmission by opioid receptor in rat subthalamic nucleus in vitro. J Physiol 541, 219–230. 10.1113/jphysiol.2001.013404.

39. Yamada, D., Takahashi, J., Iio, K., Nagase, H., and Saitoh, A. (2021). Modulation of glutamatergic synaptic transmission and neuronal excitability in the prelimbic medial prefrontal cortex via delta-opioid receptors in mice. Biochem Biophys Res Commun 560, 192–198. 10.1016/j.bbrc.2021.05.002.

40. Brenowitz, S., David, J., and Trussell, L. (1998). Enhancement of synaptic efficacy by presynaptic GABA(B) receptors. Neuron 20, 135–141. 10.1016/s0896-6273(00)80441-9.

41. Kruglikov, I., and Rudy, B. (2008). Perisomatic GABA release and thalamocortical integration onto neocortical excitatory cells are regulated by neuromodulators. Neuron 58, 911–924. 10.1016/j.neuron.2008.04.024.

42. Hjelmstad, G.O., and Fields, H.L. (2001). Kappa opioid receptor inhibition of glutamatergic transmission in the nucleus accumbens shell. J Neurophysiol 85, 1153–1158. 10.1152/jn.2001.85.3.1153.

43. Yao, Z., van Velthoven, C.T.J., Nguyen, T.N., Goldy, J., Sedeno-Cortes, A.E., Baftizadeh, F., Bertagnolli, D., Casper, T., Chiang, M., Crichton, K., et al. (2021). A taxonomy of transcriptomic cell types across the isocortex and hippocampal formation. Cell 184, 3222–3241 e3226. 10.1016/j.cell.2021.04.021.

44. Lim, L., Mi, D., Llorca, A., and Marin, O. (2018). Development and Functional Diversification of Cortical Interneurons. Neuron 100, 294–313. 10.1016/j.neuron.2018.10.009.

45. Bucurenciu, I., Kulik, A., Schwaller, B., Frotscher, M., and Jonas, P. (2008). Nanodomain coupling between Ca2+ channels and Ca2+ sensors promotes fast and efficient transmitter release at a cortical GABAergic synapse. Neuron 57, 536–545. 10.1016/j.neuron.2007.12.026.

46. Eggermann, E., and Jonas, P. (2011). How the ‘slow’ Ca(2+) buffer parvalbumin affects transmitter release in nanodomain-coupling regimes. Nat Neurosci 15, 20–22. 10.1038/nn.3002.

47. Scimemi, A., and Diamond, J.S. (2012). The number and organization of Ca2+ channels in the active zone shapes neurotransmitter release from Schaffer collateral synapses. J Neurosci 32, 18157–18176. 10.1523/JNEUROSCI.3827-12.2012.

48. Jordan, B., and Devi, L.A. (1998). Molecular mechanisms of opioid receptor signal transduction. Br J Anaesth 81, 12–19. 10.1093/bja/81.1.12.

49. Tang, K.C., and Lovinger, D.M. (2000). Role of pertussis toxin-sensitive G-proteins in synaptic transmission and plasticity at corticostriatal synapses. J Neurophysiol 83, 60–69. 10.1152/jn.2000.83.1.60.

50. Thalmann, R.H. (1988). Evidence that guanosine triphosphate (GTP)-binding proteins control a synaptic response in brain: effect of pertussis toxin and GTP gamma S on the late inhibitory postsynaptic potential of hippocampal CA3 neurons. J Neurosci 8, 4589–4602. 10.1523/JNEUROSCI.08-12-04589.1988.

51. Ikeda, S.R. (1996). Voltage-dependent modulation of N-type calcium channels by G-protein beta gamma subunits. Nature 380, 255–258. 10.1038/380255a0.

52. Aghajanian, G.K., and Wang, Y.Y. (1986). Pertussis toxin blocks the outward currents evoked by opiate and alpha 2-agonists in locus coeruleus neurons. Brain Res 371, 390–394. 10.1016/0006-8993(86)90382-3.

53. Inoue, M., Nakajima, S., and Nakajima, Y. (1988). Somatostatin induces an inward rectification in rat locus coeruleus neurones through a pertussis toxin-sensitive mechanism. J Physiol 407, 177–198. 10.1113/jphysiol.1988.sp017409.

54. Tatebayashi, H., and Ogata, N. (1992). GABAB-mediated modulation of the voltage-gated Ca2+ channels. Gen Pharmacol 23, 309–316. 10.1016/0306-3623(92)90088-2.

55. Ma, Y., Hu, H., Berrebi, A.S., Mathers, P.H., and Agmon, A. (2006). Distinct subtypes of somatostatin-containing neocortical interneurons revealed in transgenic mice. J Neurosci 26, 5069–5082. 10.1523/JNEUROSCI.0661-06.2006.

56. Naka, A., Veit, J., Shababo, B., Chance, R.K., Risso, D., Stafford, D., Snyder, B., Egladyous, A., Chu, D., Sridharan, S., et al. (2019). Complementary networks of cortical somatostatin interneurons enforce layer specific control. Elife 8. 10.7554/eLife.43696.

57. Wu, S.J., Sevier, E., Dwivedi, D., Saldi, G.A., Hairston, A., Yu, S., Abbott, L., Choi, D.H., Sherer, M., Qiu, Y., et al. (2023). Cortical somatostatin interneuron subtypes form cell-type-specific circuits. Neuron 111, 2675–2692 e2679. 10.1016/j.neuron.2023.05.032.

58. Riedemann, T. (2019). Diversity and Function of Somatostatin-Expressing Interneurons in the Cerebral Cortex. Int J Mol Sci 20. 10.3390/ijms20122952.

59. Kepecs, A., and Fishell, G. (2014). Interneuron cell types are fit to function. Nature 505, 318–326. 10.1038/nature12983.

60. Lipkin, A.M., and Bender, K.J. (2023). Axon Initial Segment GABA Inhibits Action Potential Generation throughout Periadolescent Development. J Neurosci 43, 6357–6368. 10.1523/JNEUROSCI.0605-23.2023.

61. Lovett-Barron, M., Turi, G.F., Kaifosh, P., Lee, P.H., Bolze, F., Sun, X.H., Nicoud, J.F., Zemelman, B.V., Sternson, S.M., and Losonczy, A. (2012). Regulation of neuronal input transformations by tunable dendritic inhibition. Nat Neurosci 15, 423-430, S421-423. 10.1038/nn.3024.

62. Coutens, B., and Ingram, S.L. (2023). Key differences in regulation of opioid receptors localized to presynaptic terminals compared to somas: Relevance for novel therapeutics. Neuropharmacology 226, 109408. 10.1016/j.neuropharm.2022.109408.

63. Bouchet, C.A., McPherson, K.B., Li, M.H., Traynor, J.R., and Ingram, S.L. (2021). Mice Expressing Regulators of G protein Signaling-insensitive Galphao Define Roles of mu Opioid Receptor Galphao and Galphai Subunit Coupling in Inhibition of Presynaptic GABA Release. Mol Pharmacol 100, 217–223. 10.1124/molpharm.121.000249.

64. Zhang, Z., and Pan, Z.Z. (2010). Synaptic mechanism for functional synergism between delta- and mu-opioid receptors. J Neurosci 30, 4735–4745. 10.1523/JNEUROSCI.5968-09.2010.

65. Bull, F.A., Baptista-Hon, D.T., Lambert, J.J., Walwyn, W., and Hales, T.G. (2017). Morphine activation of mu opioid receptors causes disinhibition of neurons in the ventral tegmental area mediated by beta-arrestin2 and c-Src. Sci Rep 7, 9969. 10.1038/s41598-017-10360-8.

66. Zhang, W., Yang, H.L., Song, J.J., Chen, M., Dong, Y., Lai, B., Yu, Y.G., Ma, L., and Zheng, P. (2015). DAMGO depresses inhibitory synaptic transmission via different downstream pathways of mu opioid receptors in ventral tegmental area and periaqueductal gray. Neuroscience 301, 144–154. 10.1016/j.neuroscience.2015.05.077.

67. Banghart, M.R., Neufeld, S.Q., Wong, N.C., and Sabatini, B.L. (2015). Enkephalin Disinhibits Mu Opioid Receptor-Rich Striatal Patches via Delta Opioid Receptors. Neuron 88, 1227–1239. 10.1016/j.neuron.2015.11.010.

68. Blackmer, T., Larsen, E.C., Takahashi, M., Martin, T.F., Alford, S., and Hamm, H.E. (2001). G protein betagamma subunit-mediated presynaptic inhibition: regulation of exocytotic fusion downstream of Ca2+ entry. Science 292, 293–297. 10.1126/science.1058803.

69. Capogna, M., Gahwiler, B.H., and Thompson, S.M. (1993). Mechanism of mu-opioid receptor-mediated presynaptic inhibition in the rat hippocampus in vitro. J Physiol 470, 539–558. 10.1113/jphysiol.1993.sp019874.

70. Edwards, J.G., Gibson, H.E., Jensen, T., Nugent, F., Walther, C., Blickenstaff, J., and Kauer, J.A. (2012). A novel non-CB1/TRPV1 endocannabinoid-mediated mechanism depresses excitatory synapses on hippocampal CA1 interneurons. Hippocampus 22, 209–221. 10.1002/hipo.20884.

71. Winters, B.L., Gregoriou, G.C., Kissiwaa, S.A., Wells, O.A., Medagoda, D.I., Hermes, S.M., Burford, N.T., Alt, A., Aicher, S.A., and Bagley, E.E. (2017). Endogenous opioids regulate moment-to-moment neuronal communication and excitability. Nat Commun 8, 14611. 10.1038/ncomms14611.

72. Gerachshenko, T., Blackmer, T., Yoon, E.J., Bartleson, C., Hamm, H.E., and Alford, S. (2005). Gbetagamma acts at the C terminus of SNAP-25 to mediate presynaptic inhibition. Nat Neurosci 8, 597–605. 10.1038/nn1439.

73. Hamid, E., Church, E., Wells, C.A., Zurawski, Z., Hamm, H.E., and Alford, S. (2014). Modulation of neurotransmission by GPCRs is dependent upon the microarchitecture of the primed vesicle complex. J Neurosci 34, 260–274. 10.1523/JNEUROSCI.3633-12.2014.

74. Manz, K.M., Zepeda, J.C., Zurawski, Z., Hamm, H.E., and Grueter, B.A. (2023). SNAP25 differentially contributes to G(i/o)-coupled receptor function at glutamatergic synapses in the nucleus accumbens. Front Cell Neurosci 17, 1165261. 10.3389/fncel.2023.1165261.

75. Fong, H.K., Yoshimoto, K.K., Eversole-Cire, P., and Simon, M.I. (1988). Identification of a GTP-binding protein alpha subunit that lacks an apparent ADP-ribosylation site for pertussis toxin. Proc Natl Acad Sci U S A 85, 3066–3070. 10.1073/pnas.85.9.3066.

76. Tso, P.H., Yung, L.Y., and Wong, Y.H. (2000). Regulation of adenylyl cyclase, ERK1/2, and CREB by Gz following acute and chronic activation of the delta-opioid receptor. J Neurochem 74, 1685–1693. 10.1046/j.1471-4159.2000.0741685.x.

77. Garzon, J., Rodriguez-Munoz, M., Lopez-Fando, A., and Sanchez-Blazquez, P. (2005). The RGSZ2 protein exists in a complex with mu-opioid receptors and regulates the desensitizing capacity of Gz proteins. Neuropsychopharmacology 30, 1632–1648. 10.1038/sj.npp.1300726.

78. Cruciani, R.A., Dvorkin, B., Morris, S.A., Crain, S.M., and Makman, M.H. (1993). Direct coupling of opioid receptors to both stimulatory and inhibitory guanine nucleotide-binding proteins in F-11 neuroblastoma-sensory neuron hybrid cells. Proc Natl Acad Sci U S A 90, 3019–3023. 10.1073/pnas.90.7.3019.

79. Fan, S.F., Shen, K.F., and Crain, S.M. (1991). Opioids at low concentration decrease openings of K+ channels in sensory ganglion neurons. Brain Res 558, 166–170. 10.1016/0006-8993(91)90737-g.

80. Ferre, S., Ciruela, F., Dessauer, C.W., Gonzalez-Maeso, J., Hebert, T.E., Jockers, R., Logothetis, D.E., and Pardo, L. (2022). G protein-coupled receptor-effector macromolecular membrane assemblies (GEMMAs). Pharmacol Ther 231, 107977. 10.1016/j.pharmthera.2021.107977.

81. George, S.R., Fan, T., Xie, Z., Tse, R., Tam, V., Varghese, G., and O’Dowd, B.F. (2000). Oligomerization of mu- and delta-opioid receptors. Generation of novel functional properties. J Biol Chem 275, 26128–26135. 10.1074/jbc.M000345200.

82. Waldhoer, M., Fong, J., Jones, R.M., Lunzer, M.M., Sharma, S.K., Kostenis, E., Portoghese, P.S., and Whistler, J.L. (2005). A heterodimer-selective agonist shows in vivo relevance of G protein-coupled receptor dimers. Proc Natl Acad Sci U S A 102, 9050–9055. 10.1073/pnas.0501112102.

83. Rozenfeld, R., Bushlin, I., Gomes, I., Tzavaras, N., Gupta, A., Neves, S., Battini, L., Gusella, G.L., Lachmann, A., Ma’ayan, A., et al. (2012). Receptor heteromerization expands the repertoire of cannabinoid signaling in rodent neurons. PLoS One 7, e29239. 10.1371/journal.pone.0029239.

84. Salanga, C.L., O’Hayre, M., and Handel, T. (2009). Modulation of chemokine receptor activity through dimerization and crosstalk. Cell Mol Life Sci 66, 1370–1386. 10.1007/s00018-008-8666-1.

85. Navarro, G., Cordomi, A., Casado-Anguera, V., Moreno, E., Cai, N.S., Cortes, A., Canela, E.I., Dessauer, C.W., Casado, V., Pardo, L., et al. (2018). Evidence for functional pre-coupled complexes of receptor heteromers and adenylyl cyclase. Nat Commun 9, 1242. 10.1038/s41467-018-03522-3.

86. McVey, M., Ramsay, D., Kellett, E., Rees, S., Wilson, S., Pope, A.J., and Milligan, G. (2001). Monitoring receptor oligomerization using time-resolved fluorescence resonance energy transfer and bioluminescence resonance energy transfer. The human delta -opioid receptor displays constitutive oligomerization at the cell surface, which is not regulated by receptor occupancy. J Biol Chem 276, 14092–14099. 10.1074/jbc.M008902200.

87. Moreno Delgado, D., Moller, T.C., Ster, J., Giraldo, J., Maurel, D., Rovira, X., Scholler, P., Zwier, J.M., Perroy, J., Durroux, T., et al. (2017). Pharmacological evidence for a metabotropic glutamate receptor heterodimer in neuronal cells. Elife 6. 10.7554/eLife.25233.

88. Xiang, Z., Lv, X., Lin, X., O’Brien, D.E., Altman, M.K., Lindsley, C.W., Javitch, J.A., Niswender, C.M., and Conn, P.J. (2021). Input-specific regulation of glutamatergic synaptic transmission in the medial prefrontal cortex by mGlu(2)/mGlu(4) receptor heterodimers. Sci Signal 14. 10.1126/scisignal.abd2319.

89. Zhang, L., Zhang, J.T., Hang, L., and Liu, T. (2020). Mu Opioid Receptor Heterodimers Emerge as Novel Therapeutic Targets: Recent Progress and Future Perspective. Front Pharmacol 11, 1078. 10.3389/fphar.2020.01078.

90. Yarur, H.E., Casello, S.M., Tsai, V.S., Enriquez-Traba, J., Kore, R., Wang, H., Arenivar, M., and Tejeda, H.A. (2023). Dynorphin /kappa-opioid receptor regulation of excitation-inhibition balance toggles afferent control of prefrontal cortical circuits in a pathway-specific manner. Mol Psychiatry 28, 4801–4813. 10.1038/s41380-023-02226-5.

91. Gonzalez-Burgos, G., Cho, R.Y., and Lewis, D.A. (2015). Alterations in cortical network oscillations and parvalbumin neurons in schizophrenia. Biol Psychiatry 77, 1031–1040. 10.1016/j.biopsych.2015.03.010.

92. Lewis, D.A. (2011). The chandelier neuron in schizophrenia. Dev Neurobiol 71, 118–127. 10.1002/dneu.20825.

93. Veres, J.M., Nagy, G.A., Vereczki, V.K., Andrasi, T., and Hajos, N. (2014). Strategically positioned inhibitory synapses of axo-axonic cells potently control principal neuron spiking in the basolateral amygdala. J Neurosci 34, 16194–16206. 10.1523/JNEUROSCI.2232-14.2014.

94. Zhu, Y., Stornetta, R.L., and Zhu, J.J. (2004). Chandelier cells control excessive cortical excitation: characteristics of whisker-evoked synaptic responses of layer 2/3 nonpyramidal and pyramidal neurons. J Neurosci 24, 5101–5108. 10.1523/JNEUROSCI.0544-04.2004.

95. Larkum, M.E., Wu, J., Duverdin, S.A., and Gidon, A. (2022). The Guide to Dendritic Spikes of the Mammalian Cortex In Vitro and In Vivo. Neuroscience 489, 15–33. 10.1016/j.neuroscience.2022.02.009.

96. Palmer, L.M., Schulz, J.M., Murphy, S.C., Ledergerber, D., Murayama, M., and Larkum, M.E. (2012). The cellular basis of GABA(B)-mediated interhemispheric inhibition. Science 335, 989–993. 10.1126/science.1217276.

97. Cichon, J., and Gan, W.B. (2015). Branch-specific dendritic Ca(2+) spikes cause persistent synaptic plasticity. Nature 520, 180–185. 10.1038/nature14251.

98. Xu, T.X., and Yao, W.D. (2010). D1 and D2 dopamine receptors in separate circuits cooperate to drive associative long-term potentiation in the prefrontal cortex. Proc Natl Acad Sci U S A 107, 16366–16371. 10.1073/pnas.1004108107.

99. Tang, Z., Li, S., Han, P., Yin, J., Gan, Y., Liu, Q., Wang, J., Wang, C., Li, Y., and Shi, J. (2015). Pertussis toxin reduces calcium influx to protect ischemic stroke in a middle cerebral artery occlusion model. J Neurochem 135, 998–1006. 10.1111/jnc.13359.

100. Chamberland, S., Nebet, E.R., Valero, M., Hanani, M., Egger, R., Larsen, S.B., Eyring, K.W., Buzsaki, G., and Tsien, R.W. (2023). Brief synaptic inhibition persistently interrupts firing of fast-spiking interneurons. Neuron 111, 1264–1281 e1265. 10.1016/j.neuron.2023.01.017.

101. Bender, K.J., and Trussell, L.O. (2009). Axon initial segment Ca2+ channels influence action potential generation and timing. Neuron 61, 259–271. 10.1016/j.neuron.2008.12.004.

