## Supplemental Material for "Delta opioid receptors engage multiple signaling cascades to differentially modulate prefrontal GABA release with input and target specificity"

**Supplemental Figures 1-5**

**Methods**

### SUPPLEMENTAL FIGURES

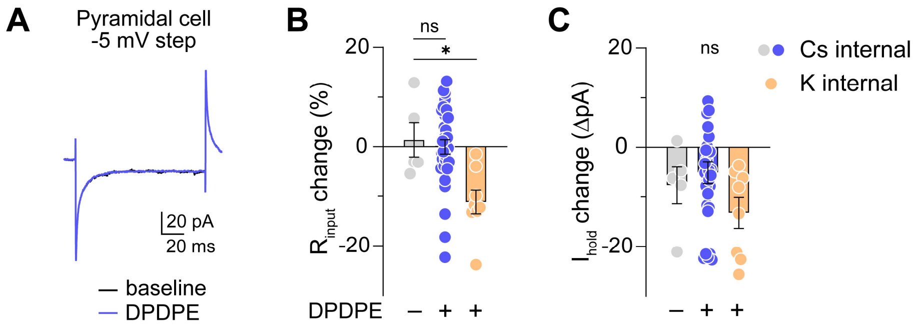


**Extended Data 1: Postsynaptic effects of DPDPE in L5 pyramidal neurons.**

(A) Example current trace from L5 pyramidal neuron in response to a -5 mV hyperpolarizing step at baseline (black) and after 1 µM DPDPE (blue).

(B) Summary data from multiple cells representing change in $R_{input}$ caused by DPDPE compared to time-locked controls (grey). Recordings using Cs-rich internal solution (blue) are compared to those using K-rich internal (peach).

(C) Same as in B but for change in holding current. * $p<0.05$.

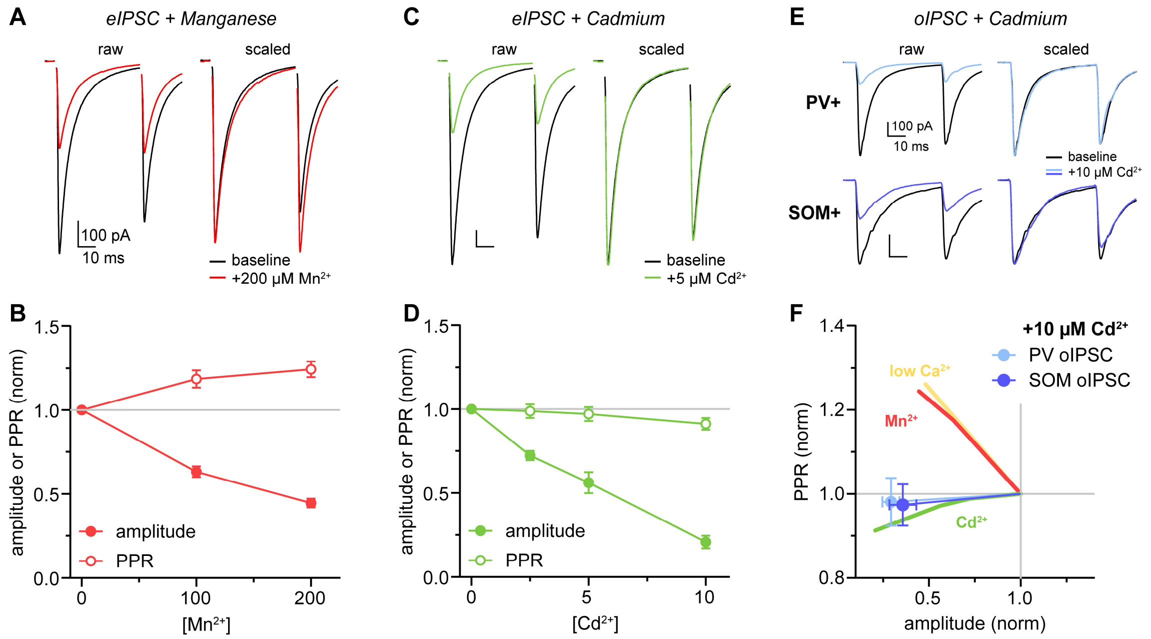


**Extended Data 2: Divalent $Ca_V$ antagonists indicate nanodomain configuration in PV+ and SOM+ terminals.**

(A) Example current traces of eIPSC pairs before and after application of manganese.
(B) Summary plot of normalized eIPSC amplitude and PPR data for multiple concentrations of manganese.
(C) Same as A but for cadmium.
(D) Same as B but for cadmium.
(E) Example traces of PV- and SOM-oIPSCs before and after cadmium.
(F) Summary normalized amplitude-PPR plot depicting slope relationships of PV- and SOM-oIPSCs with cadmium, and eIPSCs after manganese, cadmium, and low Ca.

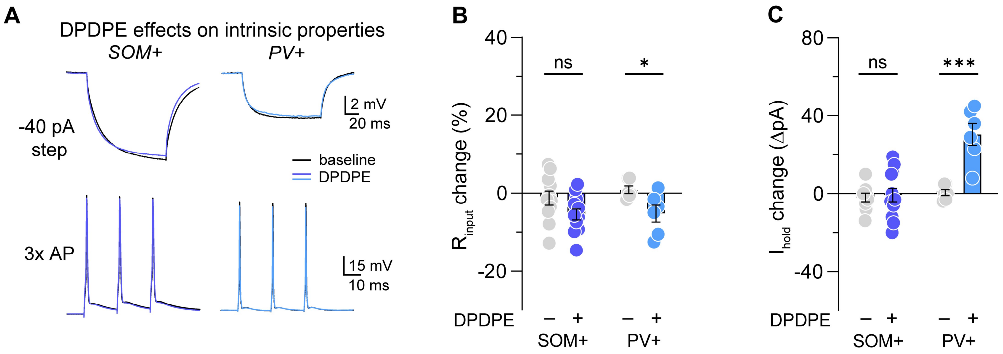


**Extended Data 3: Postsynaptic effects of DPDPE in L5 interneurons.**

(A) Example voltage traces from PV+ and SOM+ interneurons in response to a -40 pA hyperpolarizing step (top) or AP triplet stimulus (bottom) at baseline (black) and after DPDPE (blue).

(B) Summary data from multiple PV+ and SOM+ cells representing change in $R_{input}$ caused by DPDPE compared to time-locked controls.

(C) Same as in B but for change in holding current. * $p<0.05$, *** $p<0.001$.

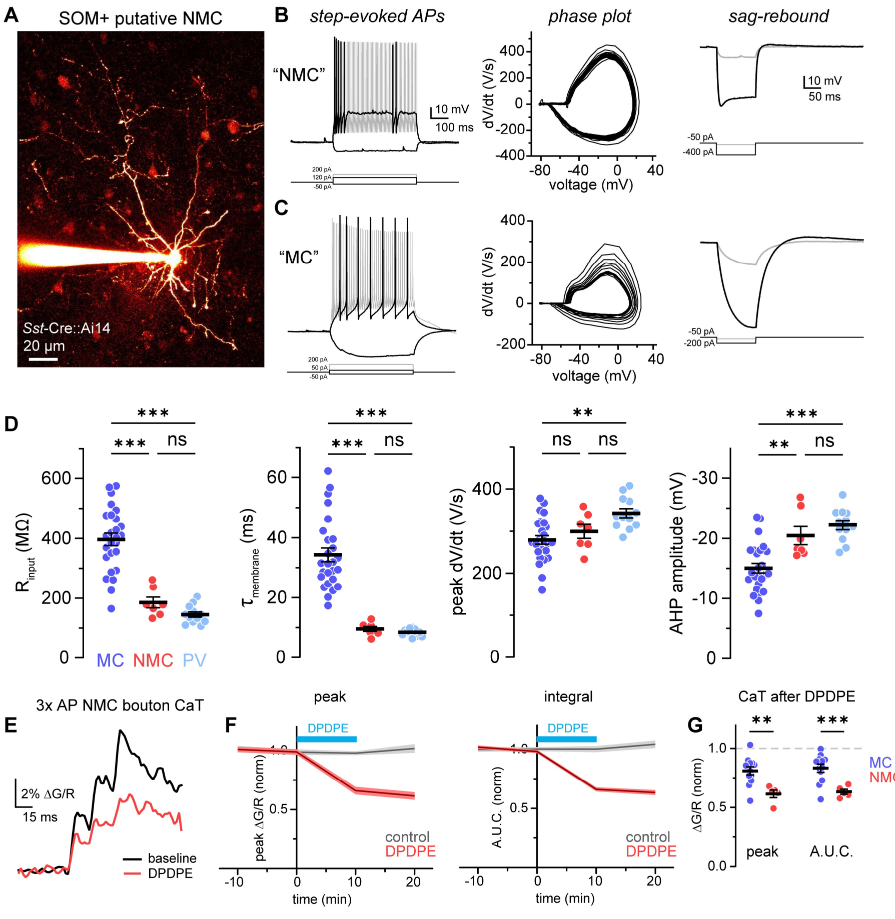


**Extended Data 4: Different neurophysiological properties of SOM+ neuronal subclasses.**

(A) Z-stack image of filled SOM+ putative non-Martinotti cell (NMC).

(B) Left: Overlaid voltage traces from an example putative NMC in response to multiple current steps depicting hyperpolarization, near-rheobase, and high frequency AP firing. Middle: Example phase-plot derived from near-rheobase AP firing. Right: Example voltage responses from weakly and strongly hyperpolarizing steps.

(C) Same as in B, but for a putative SOM+ Martinotti cell (MC).

(D) Summary plots of multiple neurophysiological measures comparing data from PV+ neurons with the two identified SOM+ subclasses.

(E) Example Ca transients evoked by AP triplet stimulus in an NMC bouton before (black) and after DPDPE (red).

(F) Summary time course plots of normalized ΔG/R peak and integral across multiple putative NMC recordings comparing time-locked vehicle control to DPDPE application.

(G) Summary normalized ΔG/R peaks and integrals (area under curve) between putative MC and NMC recordings. ** $p<0.01$, *** $p<0.001$.

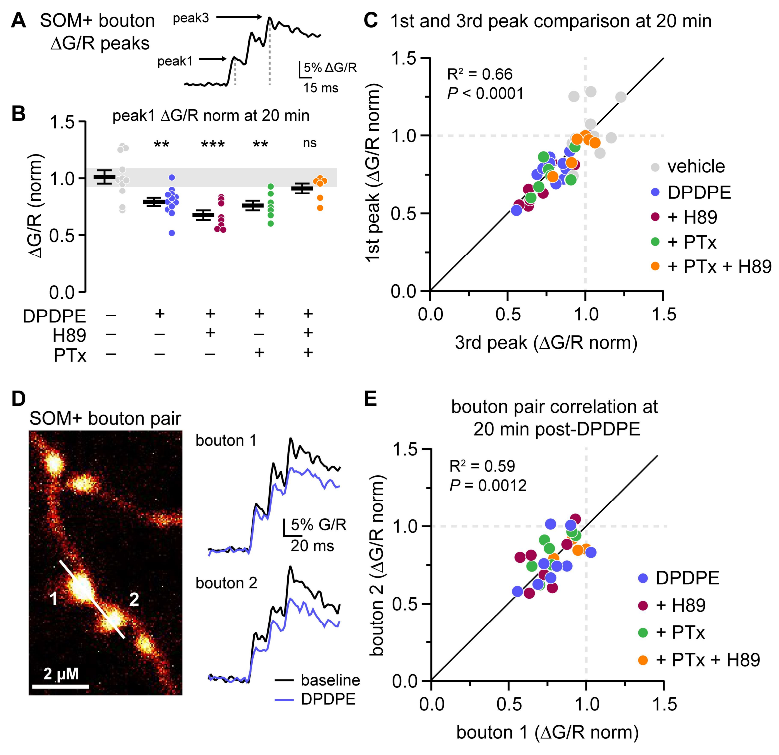


**Extended Data 5: Additional SOM+ bouton Ca transient comparisons.**

(A) Example Ca transient from SOM+ cell indicating location of ΔG/R peak measurements.
(B) Summary plot of normalized ΔG/R 1$^{st}$ peaks comparing vehicle to drug conditions. Group differences are comparable to summary 3$^{rd}$ peak data shown in Figure 4J.
(C) Plot of normalized 1$^{st}$ peak amplitudes compared to 3$^{rd}$ peak amplitudes in individual bouton recordings across drug conditions. Within-cell measurements are highly correlated.
(D) Left: Z-stack image of multiple boutons along an axonal section of a filled SOM+ neuron. White bar depicts ROI of linescan. Right: Example Ca transients from each bouton captured during linescanning before and after DPDPE application. Numbers correspond to image in left panel.
(E) Plot comparing normalized ΔG/R peaks between neighboring boutons in imaged pairs across drug conditions. ** p<0.01, ***p<0.001.

### METHODS

#### Mouse strains

All procedures were performed in accordance with UCSF IACUC guidelines. All experiments were performed on mice housed under standard conditions with *ad libitum* access to food and water. C57BL/6J wild-type (JAX: 000664), C57::PV-Cre$^{+/-}$ (JAX: 017320) and C57::SOM-Cre$^{+/-}$ (JAX: 028864) mice were used for synaptic stimulation experiments. PV-Cre$^{+/-}$::Ai14 and SOM-Cre$^{+/-}$::Ai14 mice were used for 2-photon imaging experiments in cortical interneurons. Mice aged postnatal day (P) 30-90 were used for all experiments. No significant differences based on sex were observed, and data were pooled between sexes.

#### Stereotaxic injections

Subjects were injected with virus at P30-45. Prior to viral injection, mice were anesthetized by isofluorane and head-fixed in a stereotaxic frame. For channelrhodopsin stimulation experiments, subjects were then unilaterally injected with 400-600 nL of AAV5-EF1α-DIO-hChR2(H134R)-eYFP (“DIO-ChR2”) virus into left prefrontal cortex (from bregma: A/P +1.70, M/L -0.35, D/V -2.60 mm). Acute slices were prepared approximately 4-6 weeks after injection. For PTx experiments, fresh lyophilized pertussis toxin powder (50 µg, Tocris) was obtained and reconstituted in PBS to 0.5 µg/µL and stored at 4°C (maximum 3 weeks). 2-2.5 µL PTx solution was injected into left PFC (same coordinates as above) at a rate of 150 nL/min. Following injection, the stereotaxic needle was left in place for 15 min before removing. Mice were used 24-72 hours after surgery for electrophysiology experiments.

#### Slice preparation

All experiments were performed in accordance with guidelines set by the University of California Animal Care and Use Committee. Mice aged P30-90 were anesthetized under isoflurane. Brains were dissected and placed in 4°C cutting solution consisting of (in mM): 87 NaCl, 25 $NaHCO_3$, 25 glucose, 75 sucrose, 2.5 KCl, 1.25 $NaH_2PO_4$, 0.5 $CaCl_2$, and 7 $MgCl_2$ and bubbled with 5% $CO_2$/95% $O_2$. 250 µm-thick coronal slices that included the medial prefrontal cortex were obtained via vibrating blade microtome (VT1200S, Leica). Slices were then incubated for 30 min at 33 °C in a holding chamber with artificial cerebrospinal fluid (ACSF; see below). Following 30-minute incubation, slices in holding chamber were placed at room temperature until recording, up to 8 hours. All recordings were performed in ACSF at 32-34°C.

#### *Ex vivo* electrophysiology

Slices were placed in recording chamber with circulating ACSF perfusion containing (in mM): 125 NaCl, 2.5 KCl, 1 $MgCl_2$, 25 $NaHCO_3$, 1.25 $NaH_2PO_4$, 25 glucose, bubbled with 5% $CO_2$/95% $O_2$, osmolarity adjusted to ~300 mOsm. Before recording, ACSF was supplemented with 1.3 mM $CaCl_2$ from stock solution (except for 0.65 mM Ca experiments). Neurons were identified using differential interference contrast (DIC) optics for conventional visually-guided whole-cell recording, or with 2-photon-guided imaging of tdTomato fluorescence overlaid on a scanning DIC image of the slice. Patch electrodes were pulled from Schott 8250 glass (3-4 MΩ tip resistance). For voltage-clamp IPSC recordings, patch electrodes were filled with an internal solution that contained (in mM): 80 KCl, 40 $CsMeSO_4$, 10 HEPES, 4 NaCl, 5 QX-314, 10 $Na_2$-phosphocreatine, 4 Mg-ATP, 0.4 $Na_2$-GTP; 290 mOsm, pH 7.2-7.25. For current-clamp and voltage-clamp EPSC recordings, internal solution contained (in mM): 113 K-Gluconate, 9 HEPES, 4.5 $MgCl_2$, 14 $Tris_2$-phosphocreatine, 4 $Na_2$-ATP, 0.3 Tris-GTP; 290 mOsm, pH 7.2-7.25. Each recording day (if not imaging), internal solution was supplemented with 0.1 mM EGTA from stock.

Electrophysiological recordings were collected with a Multiclamp 700B amplifier (Molecular Devices) and a custom data acquisition program in Igor Pro software (Wavemetrics). Voltage-clamp recordings of synaptic currents were acquired at 20 kHz and filtered at 3 kHz. Current-clamp recordings were acquired at 20-50 kHz and filtered at 3-20 kHz. Whole-cell compensation of pipette capacitance and series resistance ($R_s$; 50%) were applied upon patch breakthrough. All data were corrected for measured junction potentials of +8 and +12 in high

$Cl^-$ and K-gluconate-based internals, respectively. All recordings were made using a quartz electrode holder to minimize electrode drift within the slice. Data inclusion was based on the following criteria: $R_s$ <16 MΩ at baseline and <15% change over recording; holding leak current <-150 pA and <50 pA change; temperature change <1.5°C; $R_{input}$ <20% change. AP properties were measured in the first AP elicited by the current step after rheobase (rheobase+1). AP threshold was defined as the membrane potential when dV/dt first crossed 15 V/s. AHP amplitude was defined as the minimum voltage between APs relative to AP threshold. Membrane time constant ($\tau_{membrane}$) was determined by fitting an exponential to the hyperpolarization induced by a -50 pA step. $R_{input}$ was determined by -5 mV step in voltage-clamp (pyramidal), or multiple hyperpolarizing steps in current-clamp and taking the slope of the I-V relationship (interneurons).

Extracellular afferent stimulation was applied through a bipolar silver electrode in theta glass pipette connected to a battery. Patched neurons were held at -76 mV throughout recording. Pairs of stimuli were elicited through a pulse generator (0.2-0.8 V amplitude, 200 µs duration, 20 Hz frequency) at 15 s intervals to avoid induction of synaptic plasticity. The extracellular stimulating pipette was embedded in layer 5 neuropil ~100-150 µm lateral from the recorded neuron to avoid direct stimulation. Optical stimulation of ChR2-expressing afferents was made via pairs of blue light flashes (472 nm LED, 0.5-2 $mW/mm^2$, 0.1-2 ms duration, 20 Hz frequency) through the 40X objective while whole-cell recording. To avoid polysynaptic responses, voltage amplitude/LED power were tuned so that responses were synchronous and monosynaptic, no failures were observed, and E/IPSC variability between sweeps was minimized. If necessary, the stimulating electrode or LED field-of-view would be moved to recruit more stable inputs.

For PTx experiments, mPFC slices from mice locally-injected with PTx were obtained as normal. Slices near the injection site were chosen for recording. Saturating PTx effect was verified in the first slice of each recording day by bath-applying 1 µM baclofen while measuring electrically-evoked EPSCs in L5 pyramidal neurons. If EPSC amplitude changed by more than 15%, the injection was considered a failure and no slices from that animal were used for DOR experiments. Previous attempts to achieve saturating PTx concentration in PFC included intraperitoneal[1] and intracerebroventricular[2] injection of PBS-PTx solution, where 1-1.5 µg total PTx was administered to each animal. Although significant differences were observed compared to control, the effect of baclofen was reduced by a maximum of ~20%. Sufficient block (~90%) of $G_{i/o}$ signaling was only achieved through local PTx injection.

**2-photon microscopy**

For imaging experiments, a modified K-Gluconate-based internal was used where EGTA was replaced with 250 µM Fluo-5F and 20 µM Alexa 594 (Invitrogen). 2-photon laser scanning microscopy was performed as previously described[3]. A Coherent Ultra II was tuned to 810 nm for Ca imaging and morphology experiments. Fluorescence signals were captured either through a 40X, 0.8 NA objective for morphological imaging or a 60X, 1.0 NA objective for bouton Ca imaging paired with a 1.4 NA oil immersion condenser (Olympus). For Ca imaging, fluorescence was split into red and green channels using dichroic mirrors and band-pass filters (575 DCXR, ET525/70 m-2p, ET620/60 m-2p, Chroma). Green fluorescence (Fluo-5F) was captured with 10770-40 photomultiplier tubes (PMTs) selected for high quantum efficiency and low dark counts (Hamamatsu). Red fluorescence (Alexa 594) was captured with R9110 PMTs.

During bouton imaging, PV+ or SOM+ neurons were first identified using tdTomato signal overlaid on DIC scanning of PV- or SOM-Ai14 reporter mice, respectively. Neurons were then patched and neurophysiological characterization was performed with cells held near -80 mV. Boutons were identified by imaging Alexa 594 in the red channel only and searching for thin processes relatively close to the slice surface exhibiting the characteristic string-of-pearls appearance (see Figures 3E and 4F). Ca imaging was initiated >15 minutes following patch breakthrough to allow sufficient dialysis and equilibration of both Alexa and Fluo dyes in distant axonal processes. Pairs of boutons were chosen if they were in the same z-plane and could be imaged simultaneously. This allowed us to detect instances of propagation failure and also compare modulation between neighboring boutons (**Extended Data 5D,E**). Trains of 3 APs were elicited through somatic current injection (1-2 nA amplitude, 1 ms duration, 50 Hz). Fluo-5F and Alexa 594 signals were collected simultaneously in linescan mode (~600 Hz), with line ROIs encompassing both boutons. Each measurement consisted of 10 linescans repeated at 5 sec intervals. After an initial 10 scans following patch equilibration (>15 min), a linescan bout was taken every 10 min for another 30 minutes, where drug application occurred after the 2$^{nd}$ bout. Between bouts, adjustments were made at low laser power to keep both boutons in the imaging plane.

Analysis was performed on 10-linescan averages at each timepoint. ΔG/R peaks were determined by fitting an exponential to the fluorescence decay following the final stimulus. We also measured ΔG/R of the initial peak in the triplet stimulus by averaging fluorescence over a 10 ms window surrounding the first AP. These measurements were highly correlated with each other (**Extended Data 5A,C**). Furthermore, group differences in SOM+ bouton Ca transients across drug conditions were similar when using initial peaks compared to final peaks (**Extended Data 5B**). Peak extraction was performed in Matlab and smoothed with a 3-point binomial filter for presentation. Maximum intensity image projections are displayed using the "Red Hot" lookup table in FIJI. Full neuronal and dendritic reconstructions were stitched together using pairwise stitching in FIJI before generation of maximum intensity projections.

**Quantification and statistical analysis**

Data are summarized by mean ± standard error (SEM) with individual data points overlaid. Number of recordings per group are denoted in the text as n for cells and N for mice. Normality for each group was tested using Kolmorogov-Smirnov test, and group differences were assessed with parametric or non-parametric tests accordingly. Data were mostly normally distributed, and paired or unpaired t-tests were used unless otherwise specified. Statistical analysis was performed using Prism (GraphPad).

### SUPPLEMENTAL REFERENCES


1. Tang, Z., Li, S., Han, P., Yin, J., Gan, Y., Liu, Q., Wang, J., Wang, C., Li, Y., and Shi, J. (2015). Pertussis toxin reduces calcium influx to protect ischemic stroke in a middle cerebral artery occlusion model. J Neurochem *135*, 998-1006. 10.1111/jnc.13359.
2. Chamberland, S., Nebet, E.R., Valero, M., Hanani, M., Egger, R., Larsen, S.B., Eyring, K.W., Buzsaki, G., and Tsien, R.W. (2023). Brief synaptic inhibition persistently interrupts firing of fast-spiking interneurons. Neuron *111*, 1264-1281 e1265. 10.1016/j.neuron.2023.01.017.
3. Bender, K.J., and Trussell, L.O. (2009). Axon initial segment Ca2+ channels influence action potential generation and timing. Neuron *61*, 259-271. 10.1016/j.neuron.2008.12.004.
